# Programmable promoter editing for precise control of transgene expression

**DOI:** 10.1101/2024.06.19.599813

**Authors:** Sneha R. Kabaria, Yunbeen Bae, Mary E. Ehmann, Adam M. Beitz, Brittany A. Lende-Dorn, Emma L. Peterman, Kasey S. Love, Deon S. Ploessl, Kate E. Galloway

## Abstract

Subtle changes in gene expression direct cells to distinct cellular states. Identifying and controlling dose-dependent transgenes require tools for precisely titrating expression. To this end, we developed a highly modular, extensible framework called DIAL for building editable promoters that allow for fine-scale, heritable changes in transgene expression. Using DIAL, we increase expression by recombinase-mediated excision of spacers between the binding sites of a synthetic zinc finger transcription factor and the core promoter. By nesting varying numbers and lengths of spacers, DIAL generates a tunable range of unimodal setpoints from a single promoter. Through small-molecule control of transcription factors and recombinases, DIAL supports temporally defined, user-guided control of transgene expression that is extensible to additional transcription factors. Lentiviral delivery of DIAL generates multiple setpoints in primary cells and iPSCs. As promoter editing generates stable states, DIAL setpoints are heritable, facilitating mapping of transgene levels to phenotypes. The DIAL framework opens new opportunities for tailoring transgene expression and improving the predictability and performance of gene circuits across diverse applications.

**Highlights:** Promoter editing generates unimodal setpoints from a single synthetic promoter

DIAL setpoints are tuned by the length of an excisable spacer within the promoter

DIAL setpoints are uniform and robust to varying levels of the transcription factor

DIAL transmits transient inputs into heritable promoter states

The TET-DIAL system enables small-molecule activation at defined setpoints

DIAL regulates physiologically-relevant transgenes and performs in primary cells and iPSCs

**One Sentence Summary:** DIAL offers an extensible framework for designing synthetic promoters that perform across a range of cell types to generate heritable setpoints of gene expression.

## Main

Subtle changes in gene expression can generate diverging cell fates.^1–6^ Overexpression of endogenous and synthetic genes drives and redirects native processes and can augment native cellular functions.^7–10^ However, identifying which transgenes elicit these subtle effects requires fine-tuned control, and implementing control over dosage-sensitive regimes remains a challenge.^1,11^ In particular, non-linear effects of gene expression can confound inference of positive and negative regulation of phenotypes.^1,2,12,13^ Tools that support fine-scale titration of expression reveal non-monotonic relationships between expression of regulators and phenotypes.^1,2,12^ While useful for identifying linear regulators, large-scale screening tools such as CRISPR-based knockout, knockdown, and activation often do not provide sufficient resolution to find regulators with non-linear relationships to phenotypes. Such CRISPR-based screening does not predict how overexpression of transgenes influences cellular behaviors.^14^ As transgenes are increasingly used to augment cellular functions and program cell fate, there is a critical need for scalable tools to identify regulators with complex functions and define their influence on physiologically-relevant phenotypes.^14–17^

Tools that support titration may also enable tuning and control of transgene expression in therapeutic contexts for precision cell and gene therapy.^10^ To this end, synthetic biology aims to harness the power of native biology by interfacing native and synthetic gene regulatory networks. Dynamic synthetic circuits such as toggle switches, pulse generators, bandpass filters, and oscillators can dynamically control transgenes to direct cellular processes, states, and identities.^18–25^ However, rational *de novo* design of synthetic circuits remains challenging.^26–30^ Even simple inducible promoters can exhibit emergent, undesirable behaviors that impede the development of transgenic systems and gene circuits.^31–33^

Synthetic transcription factor systems offer constitutive and inducible control of gene expression.^10,12,34,35^ Synthetic transcription factors mimic the DNA-binding and transcriptional activation of native factors by modularly fusing high-affinity DNA-binding domains to strong transactivation domains (TADs). These synthetic transcription factors are often directed to binding sites upstream of a weak core promoter.^34–36^ Properties of transcription factor binding sites -- including number, distance to the core promoter, and affinity of binding -- influence the recruitment of transcriptional machinery and thus the level of transgene expression.^35,37^ Within a limited range, increasing levels of synthetic transcription factors or small-molecule inducers can increase the mean level of expression across a population of cells.^32^ However, these tools generally result in bimodal distributions of expression.^33,35,38^ Bimodality limits robust control of the entire population and may confound construction of a functional relationship between levels of expression and phenotypes. More complex circuits can linearize inducible systems to generate unimodal dose-responses at the cost of larger payloads and numbers of genetic parts, which may be difficult to translate to relevant cell types.^39^

To develop DIAL, we outlined a set of desirable features for a promoter system that controls transgene expression over dosage-sensitive regimes from a single promoter. First, we want a system capable of generating tunable setpoints of transgene expression that span physiologically-relevant ranges, supporting titration of transgene levels from a single promoter. Second, setpoints should be unimodal to ensure uniform induction and reliable control across an entire population of cells. Third, to ensure stable, homogenous output levels, setpoints should be robust to fluctuations in the levels of system components. Fourth, to map setpoint levels to phenotypes, setpoints should be recordable and heritable. Since phenotypes can emerge over longer timescales, an ideal synthetic promoter system generates stable setpoints via heritable changes that can be read at terminal timepoints. Fifth, to control induction, the system should be amenable to user-guided cues that implement reversible and irreversible changes in expression from a single promoter. Finally, for translational impact, the synthetic promoter system should be compact for delivery into primary cells.

Here we expand the precision and tunability of synthetic promoters by developing a system capable of generating multiple setpoints of transgene expression from a single promoter. Through a defined combination of inputs, the DIAL promoter system generates multiple unimodal setpoints via promoter editing (Fig 1A). Recombinase-based editing of the DIAL promoter excises a spacer between the binding sites and core promoter. Reducing the distance between the binding sites and core promoter increases transcriptional activity, shifting expression to a higher setpoint. Increasing the length of the spacer increases the setpoint range by reducing the expression from the pre-edited promoter. By nesting orthogonal recombinase sites, we constructed a nested DIAL promoter that generates four stable setpoints from a single promoter. Importantly, we demonstrate that DIAL setpoints are robust over a large range of transactivator levels. As promoter editing is genetically encoded, DIAL translates user-defined inputs into heritable setpoints. To further explore the generality of the spacer-excision architecture, we integrated the TET-ON system into the DIAL framework, generating TET-DIAL. TET-DIAL generates doxycycline-inducible setpoints, allowing compact, reversible control of setpoint induction. Importantly, for broad translation to diverse cell types, we demonstrate that DIAL can be delivered via lentivirus and generates setpoints of transgene expression in primary cells and human induced pluripotent stem cells.

**Figure 1.**
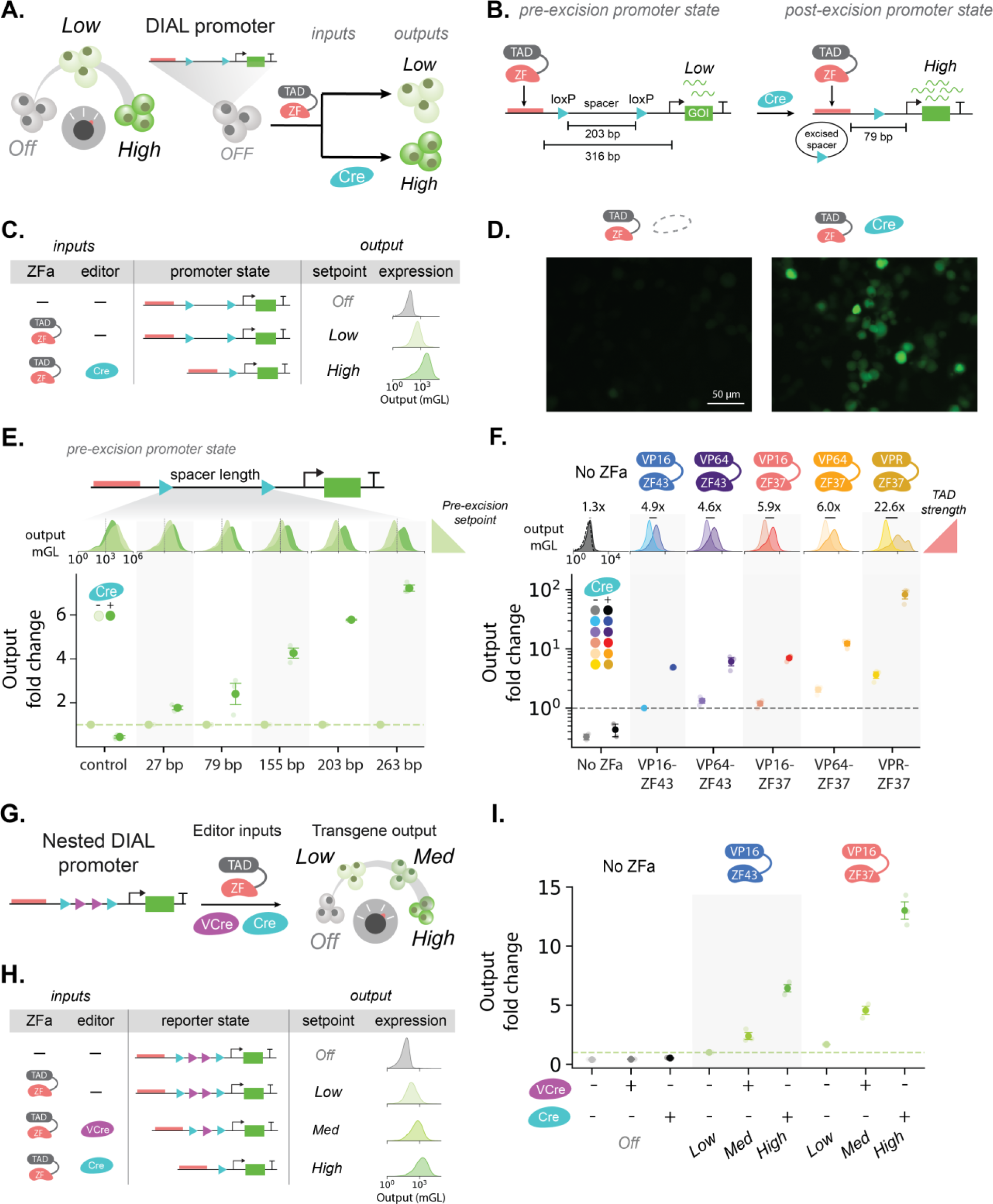
Promoter editing generates a range of unimodal setpoints from a single promoter. (A) Schematic of the DIAL promoter system. DIAL uses combinatorial inputs of synthetic zinc finger transcription factor (ZFa) and Cre recombinase to generate distinct setpoints of gene expression from a single promoter. (B) Schematic of the pre-excision and post-excision state of the 203 base pair (bp) spacer DIAL promoter before and after Cre-mediated editing, respectively. The excision of the floxed 203 bp spacer increases expression by reducing the distance between the ZF binding sites and YB_TATA minimal promoter (bent arrow) from 316 bp to 79 bp. (C) Logic table of inputs, ZFa and Cre, and outputs, expected promoter state, target setpoint, and observed gene (mGL) expression from the DIAL promoter. Output mGL single-cell distributions show output increase upon addition of ZFa (VP16-ZF37) and Cre with the 203 bp spacer DIAL promoter on plasmids intoHEK293T cells. Different combinations of inputs enables three setpoints. (D) Representative fluorescence microscopy images of mGL expressed from 203 bp spacer DIAL promoter transfected with ZFa (VP16-ZF37) on plasmids into HEK293T cells, with or without Cre. Images taken 3 days post-transfection (dpt). (E) Fold change of the output reporter expressed from DIAL promoters with different spacer lengths with co-transfected ZFa (VP16-ZF37) on plasmids into HEK293T cells, with or without Cre. Fold change is the output mGL geometric mean fluorescence intensity (MFI) normalized to the condition without Cre within each spacer length. Histograms show decreasing pre-excision expression for increasing spacer length, which generates the larger fold change upon addition of Cre. (F) Fold change of the output reporter expressed from the 203 bp spacer DIAL promoter transfected with different zinc finger activators bearing different transactivation domains (ZF-TADs, e.g. ZFa) on plasmids into HEK293T cells, with or without Cre. Fold change is the output mGL geometric MFI normalized to the condition with VP16-ZF43 without Cre. Fold change increases with stronger ZFa. (G) Schematic of nested DIAL promoter with loxP (blue) and VloxP (purple) sites. Based on the combination of ZFa, VCre, and Cre, the nested DIAL promoter generates three promoter states and four different setpoints of expression. (H) Logic table of inputs, ZFa, Cre, and VCre, and outputs, promoter state, target setpoint, and observed gene (mGL) expression for the nested DIAL promoter. Nested spacers enables four setpoints. (I) Fold change of the output reporter expressed from the nested DIAL promoter transfected on plasmids into HEK293T cells, with or without ZFa, Cre or VCre. Fold change of output mGL geometric MFI is normalized to the condition with VP16-ZF43 without either recombinase. All units for output MFI are arbitrary units (a.u.), and fold change is unitless. Large markers represent the mean of biological replicates with span indicating standard error (n=3). Histograms represent single-cell distributions sampled across bioreplicates (n=3). Statistical significance was calculated with Students t-Test with ns p>0.05; *p<0.05; **p<0.01; *** p<0.001.

## Results

### Promoter editing generates a range of unimodal setpoints from a single promoter

Native transcription factors commonly use Cys2-His2 zinc finger (ZF) binding domains to identify cognate binding sites across the genome.^40,41^ Synthetic ZF transcription factors are designed to bind arrays of unique binding sites orthogonal to native sequences.^34,35^ Zinc finger activators (ZFas) induce transcription by binding near core promoters via their ZF DNA-binding domain and recruiting transcriptional machinery via their transactivation domain (TAD). To compose the DIAL promoter system, we turned to a set of well-defined ZFas from the COMET tool kit.^34,35^ We designed the DIAL promoter with an array of tessellated binding sites specific to two ZF domains (ZF43, ZF37) separated from a downstream minimal TATA promoter by a spacer (Fig S1A-B). Between the minimal promoter and binding site array, we placed a spacer flanked by loxP (“floxed”) recognition sites. The tyrosine recombinase Cre recognizes the loxP sites and edits the promoter by excising the floxed spacer. Spacer excision brings the binding site array closer to the minimal promoter. In the presence of the ZFa, Cre-mediated excision increases expression from a low to a high setpoint (Fig 1B). In the absence of ZFa, expression is OFF. Using a combination of Cre and ZFa, we program three setpoints of gene expression from a single promoter sequence (Fig 1C).

To characterize the DIAL framework, we transfected a DIAL promoter containing a spacer length of 203 base pair (bp) into HEK293T cells. We measured fluorescence using flow cytometry. To isolate transfected cells for analysis, we gated cells based on expression of a co-transfection marker (Fig S2). Through a combination of ZFa and Cre inputs, we changed the molecular state of the 203 bp DIAL promoter to generate three unimodal output setpoints (Fig 1C-D, Fig S3A, Fig S1C). By co-expressing each ZFa with a fluorescent protein, we verified that the levels of ZFa do not change in the presence of Cre, indicating that differences in the DIAL output are not due to perturbations in levels of ZFa (Fig S3B-C).

Cre-mediated excision of the spacer increases expression by reducing the distance between the binding sites and minimal promoter. We validated spacer excision via genotyping PCR. As expected, in the presence of Cre, a shorter band appears, corresponding to the edited promoter (Fig S4). Further, addition of Cre in the presence of ZFa increases expression, reaching the level of a control promoter that lacks a spacer (Fig S1C). Next, we explored tuning the setpoints and range of the DIAL promoter. In synthetic promoter systems, expression increases as the distance between the transcription factor binding sites and the transcription start site decreases.^35^ We hypothesized that increasing the length of the excisable spacer will reduce expression of pre-excision setpoint, resulting in a larger fold-change between the low and high DIAL setpoints. For a panel of spacers ranging in length from 27 bp to 263 bp, increasing spacer length decreases pre-excision output expression and increases the fold change (Fig 1E, Fig S5A-D). Expression from the post-excision setpoint converges to the control construct which represents the post-excision sequence (Fig 1E, Fig S5C-D).

Through tessellation of binding sites, the DIAL promoter responds to ZFas with either ZF37 and ZF43 binding domains (Fig S1A), allowing us to modularly combine domains of ZFs and TADs to generate ZFas of different strengths.^35^ We found that increasing the strength of the ZFa modestly increases the setpoint and fold changes (Fig 1F, Fig S5E-F). The strongest ZFa, VPR-ZF37, substantially increases setpoint levels and the range but generates bimodal expression of the reporter, potentially limiting its utility for experiments requiring uniform control (Fig 1F). For the ZFas that generate unimodal setpoints, recombinase-mediated editing of the promoter generates a larger fold change compared to exchange of the ZFas.

To increase the number of setpoints from a single promoter, we nested a set of orthogonal recombinase sites to generate multiple excisable spacers (Fig 1G, Fig S6A-C). VCre, an orthogonal tyrosine recombinase, recognizes and excises regions flanked by VloxP sites. Addition of internal VloxP sites within the original floxed spacer allows promoter editing to generate three promoter states (Fig 1H). Addition of VCre excises the shorter, VloxP-flanked spacer, increasing the setpoint from low to medium expression. Addition of Cre excises the entire floxed spacer, inducing the high setpoint. In the absence of ZFa, DIAL expression is OFF. Through combinations of specific recombinases and ZFa, we generate four defined setpoints that span more than an order of magnitude from a single promoter construct (Fig 1I, Fig S6D-H).

### DIAL setpoints are robust to varying levels of ZFa

Cellular physiology and the process of gene regulation contribute to variability in the expression of transgenes.^42–44^ High variability in the expression of components can lead to poor performance of gene circuits.^31,32,45^ Ideally, circuits can be designed to buffer variation and ensure robust performance across populations and over time.^5,46–50^ For synthetic transcription factor systems, low expression of transcription factors contributes to bimodality, which is often masked by observation of only the mean level of the population (Fig 2A).^32^ As bimodality contributes to poor control and circuit performance, promoter systems ideally generate unimodal expression of transgenes.

**Figure 2.**
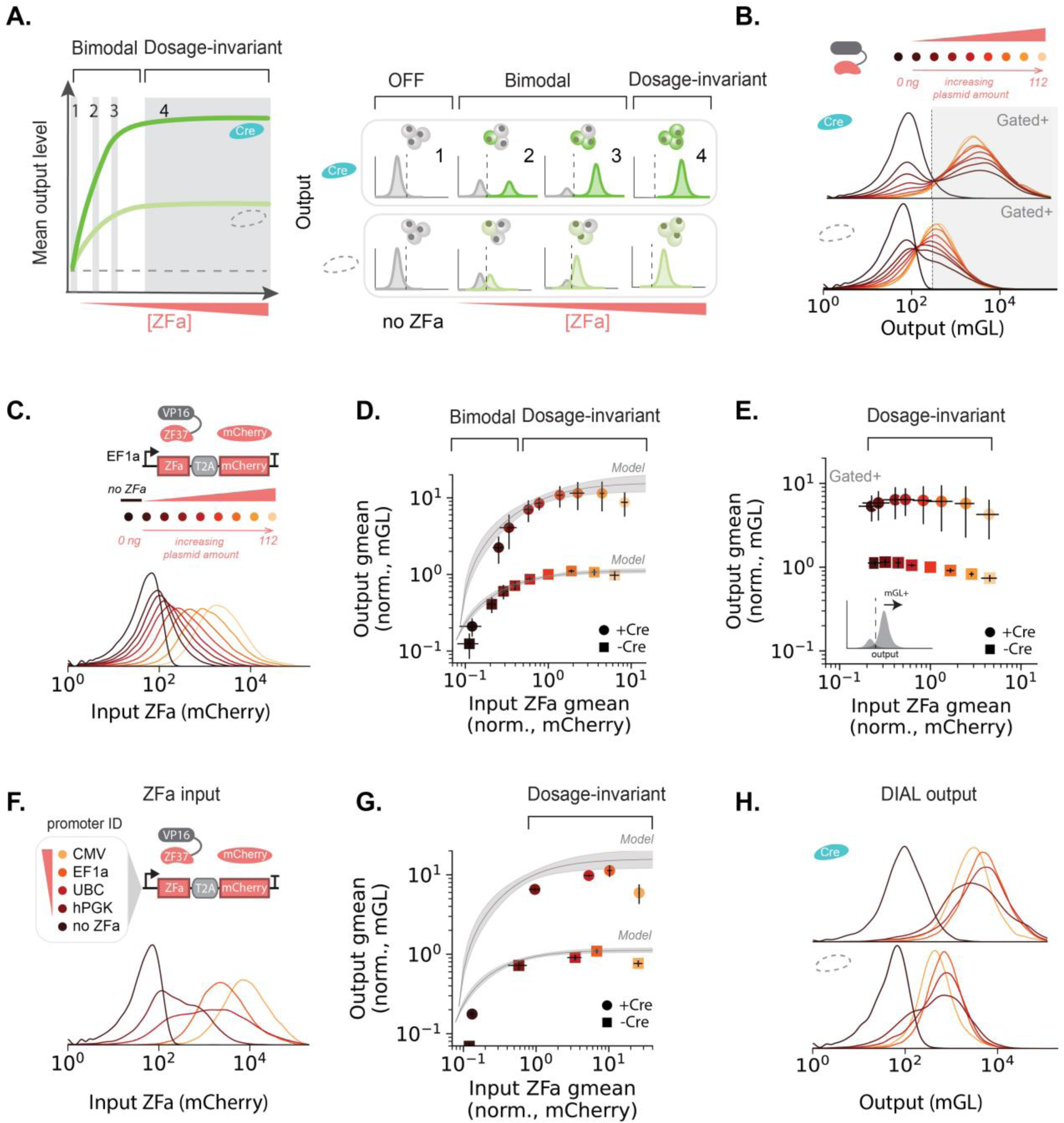
DIAL setpoints are robust to varying levels of transcription factor. (A) Mean expression from synthetic promoters increases in response to varying levels of transcription factors. Prediction of the titration of ZFa for the DIAL system. At low levels of ZFa, synthetic promoters such as DIAL generate a bimodal output, but above a threshold of ZFa level, the output levels enter a dosage-invariant regime (e.g output does not vary as ZFa varies.) (B) Single-cell distributions of output mGL expressed from the 203 bp spacer DIAL promoter titrated with ZFa (VP16-ZF37) transfected on plasmids into HEK293T cells with or without Cre. Gate drawn (mGL-A>200) to isolate populations with high output levels of the DIAL promoter. In the presence of Cre, the output mGL increases. At low levels of ZFa, output is bimodal. (C) Single-cell distributions of mCherry coexpressed with ZFa (VP16-ZF37) in B transfected on plasmids into HEK293T cells. Conditions with and without Cre are combined. Schematic of plasmid titration of ZFa. (D) Output reporter mGL geometric MFI versus input ZFa geometric MFI (proxied by co-expressed mCherry) for ZFa titration shown in B and C. Values are normalized to the condition without Cre with 0.125X (14 ng of ZFa). Overlaid lines represent model fit with 95% confidence interval. The output initially increases at low ZFa levels, where the distribution is bimodal, followed by a dosage-invariant regime. (E) Output reporter mGL geometric MFI (gated+ by the line in B) versus input ZFa geometric MFI (proxied by co-expressed mCherry, as shown in C not gated+ for mGL) transfected on plasmids into HEK293T cells with the ZFa plasmid titration. Values are normalized to the condition without Cre with 0.125X (14 ng of ZFa). Once gated, the reporter output is dosage-invariant throughout the ZFa plasmid titration. (F) Input mCherry (proxy for ZFa VP16-ZF37) single-cell distributions from constitutive promoters of varying strengths transfected on plasmids into HEK293T cells. (G) Output reporter mGL geometric MFI versus input ZFa geometric MFI (proxied by co-expressed mCherry) from different strength promoters with ZFa (VP16-ZF37) transfected on plasmids into HEK293T cells. Values are normalized to the condition without Cre with EF1a promoter. (H) Single-cell distributions of output mGL using different promoters to control ZFa transfected on plasmids into HEK293T cells, colored according to legend in F. All units for output MFI are a.u.. Large points in scatter plots represent mean of transfection biological replicates with span indicating standard error (n=6 plasmid titration, n=3 promoter strengths). Histograms represent single-cell distributions sampled across bioreplicates.

Potentially, sufficient expression of transactivators can establish regimes that are unimodal and invariant to transactivator levels.^32,33^ To examine the sensitivity of DIAL to variation in the levels of ZFa, we performed a titration of two ZFas (Fig 2B-D, Fig S7A-C). Using fluorescent markers, we measured the transgene output from the 203 bp DIAL promoter for varying levels of ZFa.

At high levels of ZFa, the mean expression from the DIAL promoter remained constant at the induced setpoint (Fig 2D, Fig S7B). In this dosage-invariant regime, the DIAL setpoints are maintained even while the level of ZFa ranges over one order of magnitude (Fig 2D, Fig S7B). At lower ZF levels, bimodality emerges. As ZF levels decrease, the overall population mean decreases due to the increasing population of cells in the OFF state (Fig 2B, D, Fig S7B, E, S8). Isolating the cells expressing the output gene, we find that DIAL expression is invariant to levels of ZFa for both the high and the low setpoints (Fig 2E, Fig S7D). At lower ZFa levels, the fraction of reporter positive cells changes but the mean of the positive cells remains unchanged (Fig S7E). While changing the levels of ZFa doesn’t uniformly change expression, DIAL provides a mechanism to uniformly increase the setpoint of expression.

To quantitatively characterize DIAL, we used a model of transcriptional activation. This model uses a simple Hill function to capture ZFa binding and the rate of transcriptional activation of the ZFa-bound promoter. Using the plasmid titration data, we fit the input ZFa and output expression for each condition in the presence or absence of Cre (Fig 2D, G, Fig S7I). Comparing the fitted parameters between the presence and absence of Cre, we could quantify the effect of promoter editing on the estimated binding affinity and rate of transcriptional activation. As expected, the binding affinity of the ZFa does not show a significant change upon excision of the spacer via Cre (Table S3-4, Fig S7J). However, decreasing the distance between the ZFa binding sites and minimal promoter increases the putative rate of transcriptional activation up to 20-fold (Table S3-4, Fig S7J).

As DIAL is robust to ZFa levels, DIAL setpoint should be flexible to the choice of promoter, opening the potential to layer control around ZFa induction. Provided ZFa levels reach the dosage-invariant regime, DIAL should generate the predicted setpoints. Using constitutive promoters of different strengths, we generated a range of ZFa expression levels (Fig 2F, Fig S7F). Changing the expression of ZFa via the selection of the ZFa promoter provided us an independent method to examine the predictions of DIAL setpoints based on ZFa expression. Consistent with the plasmid titration, output from the DIAL promoter is uniform across the range of ZFa, matching the predictions made by our model (Fig 2G-H, Fig S7G-H). Within the dosage-invariant regime, DIAL generates predictable, programmable, and highly robust unimodal setpoints, opening opportunities to layer control around system components.

### DIAL transmits transient inputs into heritable states

Transmitting transient events into heritable states supports event recording and stable direction of cell trajectories.^51–54^ As editing of the DIAL promoter is recorded in the DNA, these changes in promoter state can be transiently induced and then inherited, allowing cells to generate long-term trajectories from a temporally defined stimulus. We explored three methods of transiently inducing DIAL setpoints and tracked the heritability of DIAL when integrated (Fig 3A).

**Figure 3.**
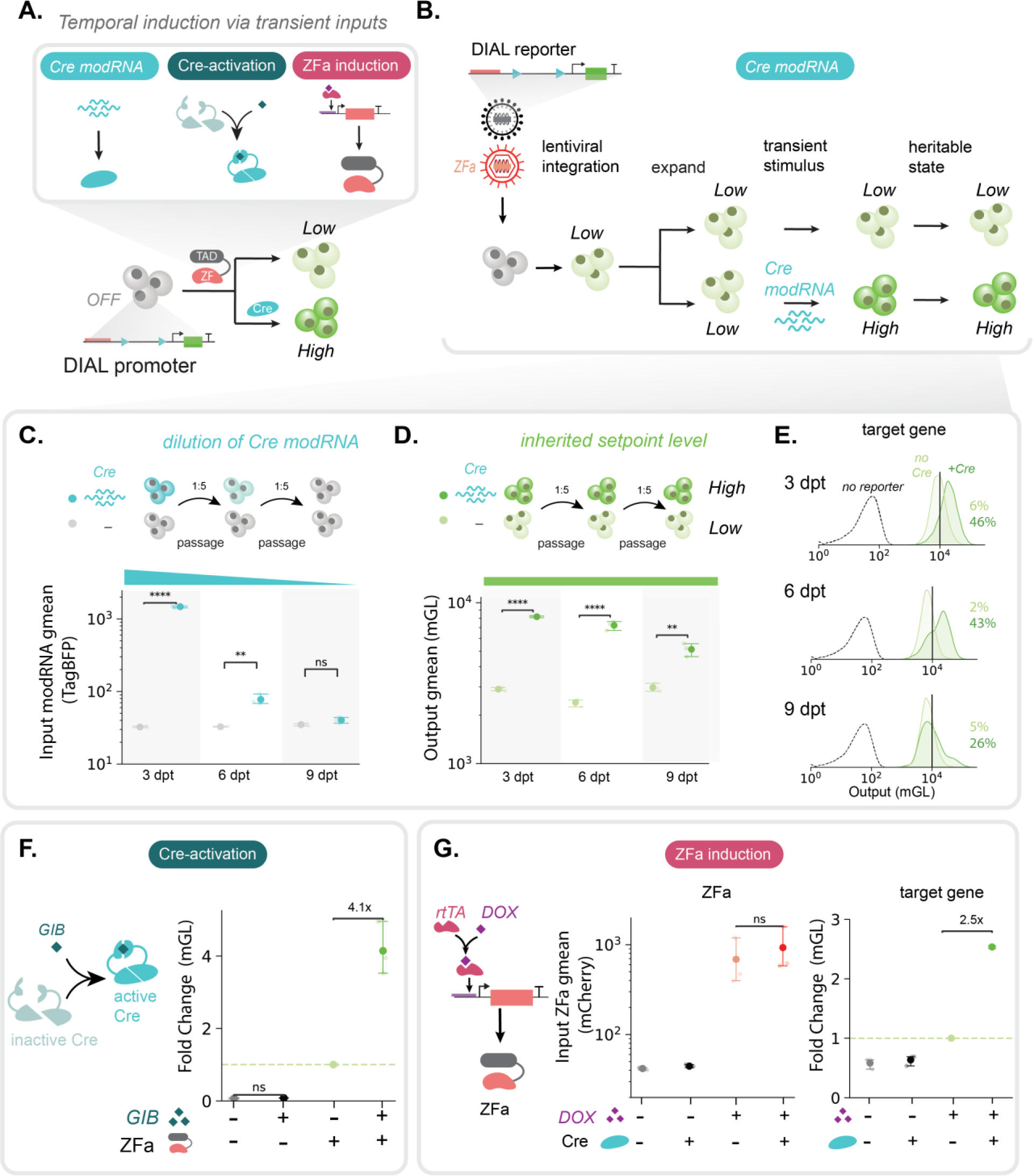
DIAL transmits transient inputs into heritable states. (A) DIAL can be regulated by different transient and temporally-defined methods to regulate the activity and expression of Cre and ZFa inputs for DIAL. (B) Steps of establishing a polyclonal HEK293T line to demonstrate heritability of DIAL setpoints. Following delivery of lentiviruses of ZFa and 203 bp spacer DIAL promoter and expansion, transfection of modRNA sets heritable setpoints of expression level (C) TagBFP (expressed via transfected modRNA, proxy for co-delivered Cre modRNA) geometric MFI over multiple passages measured on 3, 6, and 9 dpt in a polyclonal HEK293T cell line with lentivirally integrated 203-bp DIAL promoter regulating mGL and VP16-ZF37-2A-mCherry. Conditions are with and without co-transfected TagBFP modRNA and Cre modRNA added at 0 dpt. Cells are gated for mCherry+. Protein expressed from the transfected modRNA dilutes or degrades to background levels after multiple passages by 9 dpt. (D) Output reporter mGL geometric MFI from 203 bp-spacer DIAL promoter over multiple passages measured on 3, 6, and 9 dpt in a polyclonal HEK293T cell line as described in B. with. Conditions are with and without co-transfected TagBFP modRNA and Cre modRNA added at 0 dpt. Cells are gated for mCherry+. The difference in mGL setpoint level between conditions persists across multiple passages. (E) Output reporter mGL single-cell distributions over multiple passages measured on 3, 6, and 9 dpt in a polyclonal HEK293T cell line with lentivirally integrated 203 bp spacer DIAL reporter and ZFa (VP16-ZF37). Conditions are with (dark green) and without (light green) co-transfected TagBFP and Cre modRNAs at 0 dpt. Cells are gated for mCherry+. The dotted curve is a representative single-cell distribution of unstained HEK293T cells with no reporter or ZFa integrated. The vertical line represents a gate to isolate the lower peak and the higher peak. The percentages indicate the fraction of cells above the gate for the conditions with (green) and without (black) Cre modRNA. (F) Onput fold change of output mGL from the 203 bp spacer DIAL promoter transfected with gibberellin (GIB) inducible split Cre transfected on plasmids into HEK293T cells, with or without GIB (1 µM). Conditions have various combinations of co-transfected ZFa VP16-ZF37 and GIB. In the presence of GIB, activity of Cre increases which then increases expression of the output target gene, mGL, from the DIAL promoter. Fold change is the mGL geometric MFI normalized to the condition with ZFa and without GIB. (G) Input mCherry (proxy for ZFa VP16-ZF37) geometric MFI (left) and fold change of output mGL from the 203 bp spacer DIAL promoter (right) transfected with TRE3G-ZFa (shown in schematic) transfected on plasmids into HEK293T cells. Conditions have various combinations of DOX and co-transfected Cre. Presence of DOX turns output expression “OFF” or “ON”, whereas presence of Cre controls level of “ON” expression. All units for output MFI are a.u., and fold change is unitless. Points in scatter plots represent the mean of biological replicates with span indicating standard error (n=3). Histograms represent single-cell distributions sampled across bioreplicates (n=3). Statistical significance was calculated with Students t-Test with ns p>0.05; *p<0.05; **p<0.01; *** p<0.001; **** p<0.001.

To examine the heritability of DIAL states, we integrated the DIAL system into HEK293T cells using separate lentiviruses to deliver the DIAL promoter regulating mGL and the ZFa (Fig 3B). While plasmid transfection works well for delivery of recombinases in model cell lines, modRNA offers a simple method for transient *in vitro* and *in vivo* delivery of recombinases to diverse cell types.^55–57^ Delivery of modRNA encoding the Cre recombinase increases expression from DIAL in a fraction of the cells (Fig S9A). To estimate dilution of Cre over serial passaging, we co-delivered modRNAs encoding Cre and a fluorescent protein (TagBFP) (Fig 3C). By 9 days post-transfection (dpt), TagBFP levels dropped to background levels. At 9 dpt, we observe a clear separation between the background and the populations with high and low setpoints, indicating the DIAL promoter state is heritable over this period (Fig 3D, Fig S9B). However, the mean expression of the population that received modRNA Cre decreases with time, which likely reflects the expansion of unedited cells and not transgene silencing.^31^ Higher expression of transgenes can induce cellular burden and reduce proliferation.^26,28–30^ If unedited cells expand in the condition that received Cre, we would expect an increase in the fraction of cells at the lower peak, matching the expression level of the unedited cells. Conversely, transgene silencing would broadly reduce expression of the setpoints. The increase in the fraction of the population at the low setpoint suggests the loss of cells at the high setpoint reflects the combined effects of fractional editing and transgene burden (Fig 3E). The level of the higher peak does not substantially change over the 9 dpt, supporting heritability of the setpoint from the edited promoter state. Together, these data suggest that DIAL can induce heritable expression changes that can be tracked over multiple rounds of cell division. Importantly, DIAL generates a dose-dependent selection effect, which suggests that DIAL may support identification of transgenes with subtle effects on proliferation.

For genetically-engineered model organisms and *in vivo* applications where stable integration of reporter and recombinase expression may be desirable, small-molecule control of Cre offers a simple method for adjusting the DIAL setpoint. Cre-mediated editing can be induced via small-molecules and light.^58^ Addition of gibberellin (GIB) induces Cre activity via chemical-inducible dimerization (CID) domains tethered to halves of the split Cre recombinase.^58^ In transfection, addition of GIB in the presence of ZFa increased expression of the reporter (Fig 3F, Fig S9C). Small-molecule control of recombinases can also be combined or exchanged for small-molecule control of the ZFa.

DIAL relies on expression of the ZFa. Thus, inducible, pathway-responsive, and cell-type specific promoters that induce ZFa expression provide an additional method to selectively activate DIAL. Small-molecule induced expression of the ZFa provides a reversible mechanism to induce expression at the setpoint defined by the promoter state. To demonstrate inducible control of DIAL, we encoded the ZFa under the DOX-inducible TRE3G promoter. As expected, combinations of DOX and Cre generate three setpoints of expression (Fig 3G, Fig S9D-E). Unlike promoter editing with Cre, small-molecule induction of ZFa does not provide inherited memory via editing. Removal of DOX will return expression to the OFF setpoint. Altogether, DIAL setpoints can be changed and induced at user-defined timepoints through transient stimuli.

### Integration of the TET-ON system into the DIAL framework enables small-molecule activation at defined setpoints

We hypothesized that the DIAL framework could be expanded to other transactivator systems that use upstream binding sites to recruit transcriptional machinery to a core promoter. To test this hypothesis, we integrated the commonly used TET-ON system into the DIAL framework. In the presence of doxycycline (DOX), the transactivator rtTA binds to the tet-responsive promoter composed of an array of tetO sites upstream of a minimal promoter, resulting in the expression of the downstream gene. Thus, the TET-DIAL system enables reversible, small molecule-based induction of transgenes.

We hypothesized that incorporating the TET-ON system into the DIAL framework would shift expression between unimodal setpoints while maintaining reversible, small molecule-responsive induction. We constructed TET-DIAL by inserting a floxed spacer in between the tetO sites and the minimal CMV promoter (Fig 4A-B). Across a range of spacer lengths, addition of Cre increases output expression up to five-fold over pre-excision levels (Fig 4C-E, Fig S10A-D). The post-excision setpoint matches the level of expression of a control lacking a spacer, and we confirmed excision via PCR (Fig S10D).

**Figure 4.**
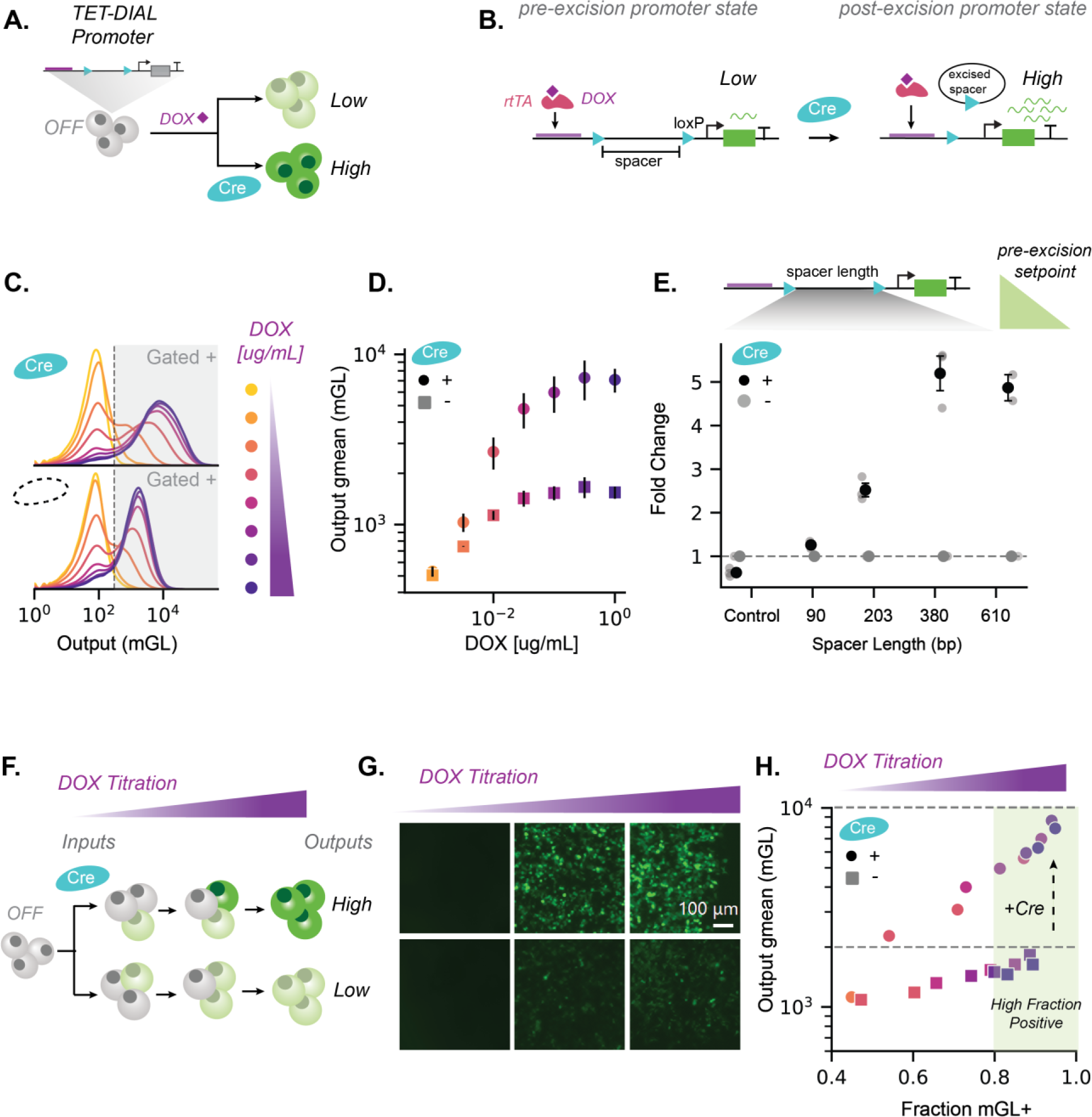
DIAL framework extends tunable setpoints across transactivator systems. (A) Schematic of TET-DIAL promoter system. With co-delivered with rtTA, TET-DIAL uses combinatorial inputs of DOX and Cre to generate distinct setpoints of gene expression from a single promoter. (B) Schematic of the pre-excision and post-excision state of the TET-DIAL promoter before and after Cre-mediated editing, respectively. The excision of the floxed spacer reduces the distance between the tetO sites and minCMV minimal promoter. (C) Output mGL single-cell distributions from the 610 bp spacer TET-DIAL promoter with rtTA transfected on plasmids into HEK293T cells, with and without Cre. DOX titration (1 ug/mL titrated down) matches colors in D. Addition of Cre increases reporter expression, whereas titrating DOX results in concurrent changes in fraction of reporter positive cells and expression level. Gate is drawn to isolate cells with expression above the no DOX condition (lightest yellow). (D) Output mGL geometric MFI versus DOX concentration for DOX titration shown in C. (E) Fold changes of the output reporter mGL expressed from the TET-DIAL promoter of varying spacer lengths transfected with rtTA on plasmids into HEK293T cells with DOX (1 µg/mL). Fold change the output mGL geometric MFI normalized to the condition without Cre within each spacer length. Longer spacer lengths generate larger fold changes upon addition of Cre. (F) Titrating DOX in the TET system leads to a concurrent increase in fraction of reporter positive cells and mean expression level. (G) Representative fluorescence microscopy images of output mGL expression from 203 bp spacer TET-DIAL promoter co-transfected with rtTA on plasmids into HEK293T cells, with or without Cre at 0, 0.3 and 1 µg/mL DOX concentrations. Images taken at 3 dpt. Scale bar, 20 µm. (H) Output mGL geometric MFI of the gated positive population versus the fraction of cells positive gated positive from the 610 bp-spacer TET-DIAL co-transfected with ZFa on plasmids into HEK293T cells, with or without Cre. Gate drawn (mGL-A>200) to isolate populations with output fluorescence above condition without DOX. DOX concentrations match the corresponding colors in (D). Dashed lines separate low and high reporter expression regimes. TET-DIAL allows for a shift between a low and high reporter expression state via promoter editing without substantial effect on the fraction of mGL positive cells. To set unimodal expression levels (off, low, and high) with high fraction positive in the “ON” state, presence of DOX at high concentration can control whether expression is “OFF” or “ON”, and presence of Cre can control the level of the “ON” expression. All units for output MFI are a.u., and fold change is unitless. Points in scatter plot D and summary plots represent mean of biological replicates with span indicating standard error(n=3). Points in scatter plot H represents individual bioreplicates (n=3). Histograms represent single-cell distributions sampled across bioreplicates (n=3).

Similar to DIAL, we observed that the output profile of TET-DIAL generates two distinct regimes of modality. At low levels of DOX, TET-DIAL generates a bimodal output whereas at higher levels of DOX the output is unimodal and insensitive to changes in DOX (Fig 4C-D, Fig S10A-B). Addition of Cre increases expression across DOX concentrations. As with ZFas, titrating the levels of the rtTA generated bimodal expression (Fig S11). Increasing DOX concentration concurrently changes the mean expression and the fraction of the reporter expressing cells (Fig 4F-H, Fig S10C). Alternatively, Cre editing of the TET-DIAL promoter allows us to independently change the setpoint without substantially changing the fraction of induced cells. Together, TET-DIAL enables unimodal shifts in output setpoint via promoter editing while retaining the reversibility and temporal control offered by small-molecule induction. Expanding the DIAL system to additional transactivator systems suggests that the DIAL framework serves as an extensible framework for building robust setpoints from a single promoter construct.

### DIAL is portable to primary cells and iPSCs and regulates diverse transgenes

For the broadest impact across research and therapeutics, genetic control systems need to perform in primary cells and human iPSCs. To characterize DIAL performance in primary cells, we delivered DIAL promoters via lentivirus into mouse embryonic fibroblasts (MEFs) (Fig 5A). Combinations of ZFa and Cre were delivered via retroviruses. As expected, the addition of Cre increases the DIAL setpoint without affecting ZFa expression (Fig 5B-C, Fig S12A-H). The longer spacer and the stronger ZFa generate a larger range between setpoints (Fig 5B, Fig S12A, C). For temporal control of output expression in MEFs, we encoded the ZFa under the control of a DOX-inducible TRE3G promoter on a lentivirus that constitutively expresses the rtTA transactivator (Fig 5D). We delivered the DIAL promoter on a separate lentivirus. As expected, delivery of Cre via retrovirus increases expression without affecting ZFa levels (Fig 5D, Fig S12G). Addition of DOX induces expression of the ZFa, activating expression from DIAL at low and high setpoints based on the presence of Cre (Fig 5D, Fig S12G).

**Figure 5.**
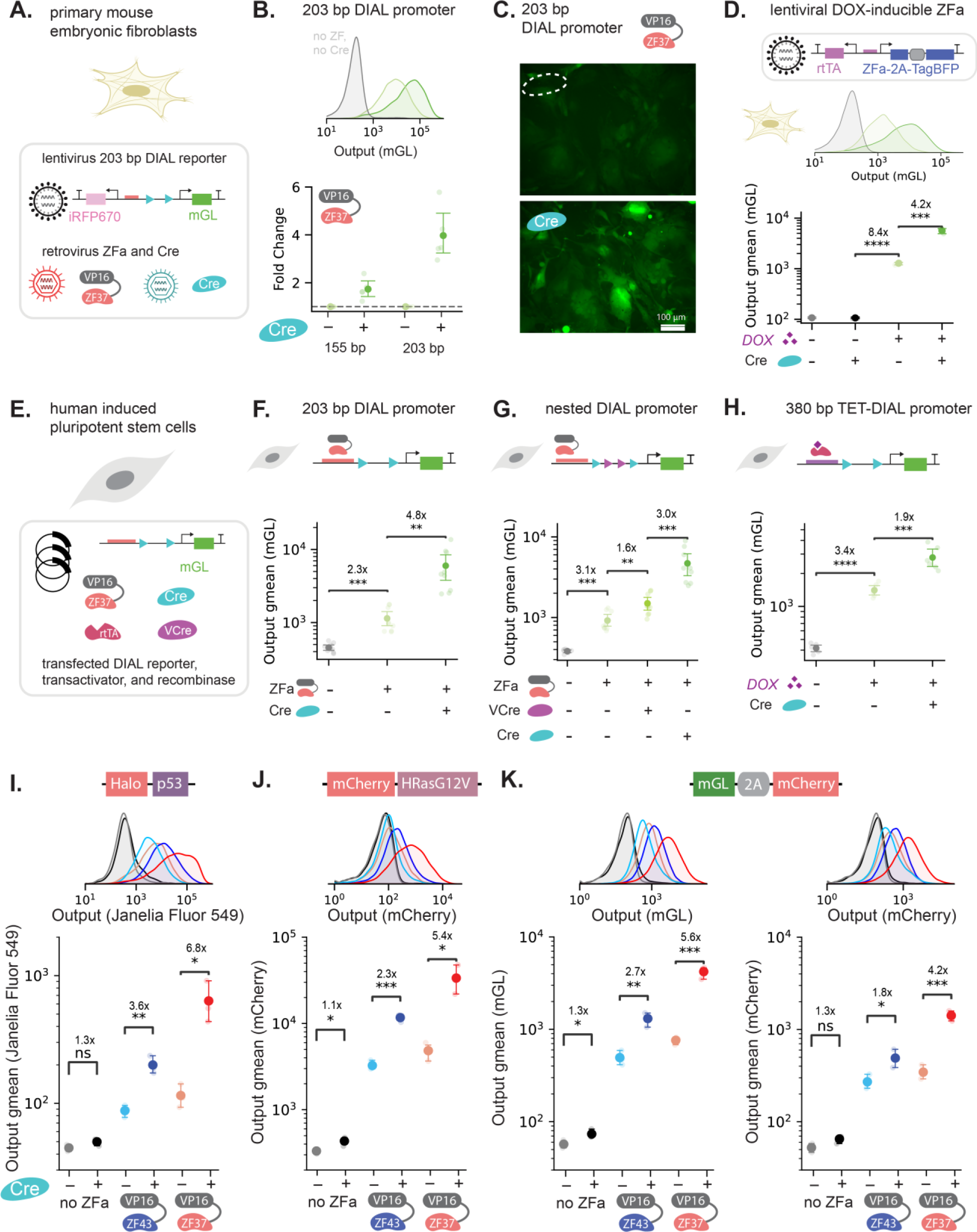
DIAL is portable to primary cells and iPSCs, and regulates diverse transgenes. (A) Delivery of the DIAL promoter system via lentivirus to primary mouse embryonic fibroblasts (MEFs) (A-C). The lentivirus encodes the DIAL promoter regulating mGL and a divergent EF1a-iRFP670. Retroviruses constitutively express Cre, VP16-ZF37-2A-mCherry, and VP16-ZF43-2A-TagBFP, and can be delivered in various combinations to set expression from the DIAL promoter. (B) Single-cell distributions of output mGL (top) expressed from 203 bp spacer DIAL promoters infected via lentivirus into MEFs, with ZFa retrovirus VP16-ZF37 and with or without Cre retrovirus. To control for infection, cells are gated by iRFP670 and mCherry when ZFa retrovirus is present. The grey curve represents cells with the DIAL promoter but without ZFa. Fold change of the output reporter (bottom) expressed from DIAL promoters with different spacers infected with ZFa via lentivirus into MEFs (gated for iRFP670+ and mCherry+). Fold change is the output mGL geometric mean fluorescence intensity (MFI) normalized to the condition without Cre within each spacer length. In the presence of ZFa, fold change is larger for longer spacer length. (C) Representative fluorescence microscopy images of mGL expressed from 203 bp spacer DIAL promoter infected via lentivirus with ZFa VP16-ZF37 retrovirus into MEFs, with or without Cre retrovirus. Images taken 3 days post-infection (dpi). (D) Single-cell distributions of output mGL (top) expressed from 203 bp-spacer DIAL promoter infected via lentivirus with lentivirus of divergent rtTA and TRE3G-ZFa via lentivirus into MEFs, with or without DOX and Cre retrovirus. Output mGL geometric MFI (bottom). Presence of DOX turns on expression, and addition of Cre unimodally increases expression from low to high. Cells are gated for iRFP670+. In the presence of DOX, cells are additionally gated for TagBFP+ (E) Schematic of various combinations of ZFa VP16-ZF37, VCre, Cre, rtTA, and DIAL promoter transfected on plasmids into iPSCs. (F) Output reporter mGL geometric MFI expressed from 203 bp spacer DIAL promoter transfected with ZFa on plasmids into iPSCs, with or without Cre. Output expression turns on in the presence of ZFa and increases upon addition of Cre. (G) Output reporter mGL geometric MFI from nested DIAL promoter in transfected on plasmids into iPSCs, with or without VP16-ZF37, VCre and Cre. Output expression turns on in the presence of ZFa and sequentially increases upon addition of VCre and Cre. (H) Output reporter mGL geometric MFI from 380 bp spacer TET-DIAL promoter transfected with rtTA on plasmids into iPSCs, with or without DOX (1 ug/mL) and Cre. Output expression turns on in the presence of DOX and increases upon addition of Cre. (I, J, K) Output reporter geometric MFI from 203 bp spacer DIAL promoter regulating different target genes co-transfected with ZFa on plasmids into HEK293T cells, with or without Cre. DifAcross different target genes, output expression turns on in the presence of ZFa and increases upon addition of Cre. All units for output MFI are arbitrary units (a.u.), and fold change is unitless.Large plot points represent mean of biological replicates (B, D, I-K; n=3) or each well in biological replicates with span indicating standard error (F-H; n=2 in H, all others n=3). Histograms represent single-cell distributions sampled across bioreplicates (n=3). Annotated fold change indicates mean of fold changes calculated within each bioreplicate. Statistical significance of bioreplicates (D, I-K) or all wells across bioreplicates (F-H) was calculated with Students t-Test with ns p>0.05; *p<0.05; **p<0.01; *** p<0.001; **** p<0.0001.

Next, we tested a range of DIAL promoters in human induced pluripotent stem cells (iPSCs) (Fig 5E). We transfected the 203 bp DIAL promoter, the nested DIAL promoter, and the 380 bp TET-DIAL promoter into iPSCs with combinations of ZFas and recombinases. As expected, addition of ZFa induces expression, and recombinase-mediated editing increases the setpoint (Fig 5F-G, Fig S13A-E). The nested DIAL promoter generates four setpoints of expression (Fig 5G, Fig S13B). For the 380 bp TET-DIAL promoter, addition of DOX induces expression and recombinase-mediated editing increases the setpoint (Fig 5H, Fig S13C). Thus, DIAL provides a toolkit that performs across a range of cell types.

Genetic controllers should be able to regulate arbitrary genes to execute titrations, enable screening, and enact diverse control functions. To demonstrate that DIAL can control expression of functionally relevant genes, we encoded the tumor suppressor protein p53, encoded by the*Trp53* gene, and HRas^G12V^, an oncogenic mutant of the *HRAS* gene, under the control of the DIAL promoter. To visualize expression and measure setpoints, we fused p53 to Halo, a ligand controlled self-labeling protein tag, and HRas^G12V^ to the fluorescent protein mCherry. As expected, DIAL generates unimodal setpoints of expression of the fusion proteins Halo-p53 and mCherry-HRasG12V without affecting ZFa level (Fig 5I-J, Fig S14). Further, DIAL regulates multiple genes by controlling expression of polycistronic cassettes (Fig 5K, Fig S14). In summary, DIAL generates setpoints of expression for arbitrary genes and in diverse cell types, supporting both research and translational applications.

## Discussion

In this work, we present DIAL, a highly modular and extensible framework for engineering synthetic promoter systems. DIAL generates multiple unimodal setpoints of expression from a single promoter sequence (Fig 1, Fig 2). Using the DIAL framework, we generate a toolkit of promoters responsive to diverse synthetic transcription factors with rationally tunable setpoints and ranges (Fig 1, Fig 4). We demonstrate that recombinase-mediated editing of the DIAL promoter increases the setpoint of the target gene. DIAL can translate transient, user-defined inputs into heritable changes in the setpoint, supporting flexibility on the timing of setpoint changes (Fig 3). For broad application, we show that DIAL can regulate physiologically-relevant transgenes and performs across diverse cell types including human iPSC and primary murine cells (Fig 5).

Precise programming of gene expression levels represents one of the most important and challenging goals in synthetic biology.^59–62^ While tools for control in mammalian systems have expanded over the last decade, few simple circuits have been translated clinically.^61^ In part, these limitations are driven by poor performance in primary cells and emergent behaviors such as bimodality and transgene silencing.^31,32,45^ We demonstrate that titration of synthetic transcription factors changes the fraction of cells in the “ON” population but does not change the mean expression of the “ON” population (Fig 2). Alternatively, promoter editing via DIAL generates distinct unimodal setpoints of expression. DIAL setpoints are robust to variation of the synthetic transcription factor (Fig 2). Over a ten-fold range of transcription factor expression, DIAL output remains constant. In the TET-DIAL system, small-molecule titration of the doxycycline-inducible system slightly tunes mean, but comes at the expense of changing the fraction of cells that express the output gene (Fig 4). Via promoter editing, DIAL uncouples setpoints and bimodality, supporting independent tuning of the induced fraction of the population and the mean levels of expression. In the dosage-invariant regime, DIAL offers unimodal expression at programmable setpoints. The combination of unimodality and dosage-invariance to component variation indicates that DIAL will improve the predictability and performance of gene circuits.

The DIAL framework allows integration of small molecule-regulated synthetic transcription factors, indicating extensibility beyond ZF-based transcription factor. With TET-DIAL, we demonstrate the DIAL framework can be extended to the doxycycline-inducible system, supporting reversible, small molecule-responsive induction (Fig 4). As the field has developed an array of synthetic promoter systems^10,63^, we envision that the DIAL framework could guide design of multiple unimodal setpoints in these systems. Similarly, integration of arrays of binding sites for diverse endogenous transcription factors into the DIAL framework may allow construction of tunable, cell-state responsive promoters.^36^ Encoding the binding sites derived from pathway-responsive and tissue-specific promoters into DIAL may allow tuning of activity from these promoters and further support circuits for information processing, recording, and computation of native signals.^25,64^ Our toolkit uses two different core promoter sequences. Exchanging these sequences for other core promoters of different types may further tune the output and range from DIAL promoters.^65,66^ We observed that spacer length translated to output ranges that varied between the DIAL and TET-DIAL, suggesting that trends in spacer lengths may be preserved but the absolute length for optimal, desired range may vary by other system features such as the synthetic transcription factor and core promoter. As contact between promoters and enhancers provides one mode of native gene regulation, it is interesting to speculate how DIAL might be used to study transcriptional dynamics.^67^ Potentially, decreasing the distance between the binding sites and core promoter increases the frequency of transcriptional bursts, as has been observed for forced enhancer looping and other synthetic transcription factor systems.^68,69^ As a well-defined system, DIAL provides a modular tool for parsing properties of gene regulation.

While recombinase systems have been used extensively for inducing expression, these systems primarily control binary states of expression (ON/OFF).^52,58,78–80^ Other systems such SCRaMbLE and GEMbLER have used recombinases to randomly recombine elements to generate diversity of expression for screening.^81–83^ In yeast, GEMbLER supports diversification of promoter sequences and expression profiles through random recombination of arrays of floxed endogenous promoters, leading to diverse expression levels.^84^ While such promoter recombination supports the identification of novel promoter combinations and optimal levels, random recombination does not support precise programming. Thus, DIAL’s ability to provide fine-scale tuning offers a unique application for tyrosine recombinases.

As more recombinases are being discovered, characterized, and engineered^70,71^, we expect that DIAL will expand to take advantage of new recombinase properties and toolkits. Nesting recombinase sites within the spacer allowed us to access more expression levels from a single DIAL promoter (Fig 1). Additional orthogonal recombinase sites could be nested to expand the number of DIAL setpoints. Orthogonal sets of recombinases could also mediate independent control of setpoints for individual genes. Given the ubiquity of recombinase tools in model organisms and cell lines, DIAL may integrate with diverse recombinase-based tools for lineage tracing.^72–76^ We show that Cre activity can be instigated by orthogonal control systems, including external user-defined cues like modRNA and small molecules, opening opportunities for changing setpoints at a specific timepoint.^58,77^

While the DIAL framework provides many insights on engineering synthetic promoter systems, there remain limitations and opportunities to expand the capabilities. While DIAL setpoints are robust to synthetic transactivator level (Fig 2, Fig S11), other factors, such as the dosage of the DIAL promoter construct itself, may influence the setpoint. Alternative types of genetic controllers may provide control for variation in DNA dosage.^50,59,60,85–87^ In its current form, DIAL allows us to increase but not decrease the level of the setpoint, limiting application to systems where a reduction in the setpoint is needed. Although permanent DNA-level editing of DIAL offers stability and long-term, hereditable memory, setpoint changes are irreversible. TET-DIAL partially overcomes this limitation. Removal of DOX allows cells to return to the OFF state, but induction requires constant delivery of the small-molecule activator and edited promoter states remain irreversible. While many applications require fine-scale changes of transgene expression, a lack of large ranges from DIAL may be a limitation for some applications. Limited range may be a result of other DIAL features. In the current form of DIAL, we selected spacer sequences which we rationally identified as putatively neutral. A more expansive characterization may identify sequence requirements for optimal performance of DIAL promoters.

Overall, DIAL expands the mammalian cell engineering toolkit to achieve precisely programmable profiles of gene expression. With ZFa and TET binding sites, we offer a set of DIAL promoters with a variety of fold changes and the ability to set multiple unimodal setpoints from a single construct. Control via a single promoter offers the scalability needed generate multiple setpoints from libraries of transgenes and inherently controls for bias of clonal founder lines for a single transgene. Thus, DIAL may help identify and control transgenes with subtle dose-dependent effects on cellular states. Finally, we anticipate that the extensible framework offered by DIAL will support expansion to novel synthetic promoter systems and broadly improve precision cell engineering.

## ACKNOWLEDGEMENTS

Research reported in this manuscript was supported by the National Institute of General Medical Sciences of the National Institutes of Health under award number R35-GM143033, by the National Science Foundation under the NSF-CAREER under award number 2339986, and the with funding from Institute for Collaborative Biotechnologies. S.R.K., B.A.L., and E.L.P. are supported by the National Science Foundation Graduate Research Fellowship Program under grant No. 1745302. A.M.B. is supported by the National Cancer Institute under award number F99CA284280. We thank the Koch Institute’s RA. Swanson (1969) Biotechnology Center (National Cancer Institute Grant P30-CA14051) for technical support, specifically the Flow Cytometry Core Facility. We thank Christopher Johnstone, Joji Teves, Diya Godavarti, and Jane Atkinson for feedback on the development of the manuscript.

## AUTHOR CONTRIBUTIONS

S.R.K., Y.B., and K.E.G. conceived and outlined the project. S.R.K., Y.B., M.E.E., and K.E.G. and analyzed DIAL and TET-DIAL characterization. A.M.B. and B.A.L. performed lentiviral delivery to primary mouse embryonic fibroblasts and analyzed data. E.L.P. and K.S.L. performed iPSC transfections. D.S.P. assisted with experimental design and production of modRNA. S.R.K., Y.B., M.E.E., A.M.B. and B.A.L., and K.E.G. wrote the manuscript. K.E.G. supervised the project.

## DATA AND MATERIALS AVAILABILITY

Data needed to evaluate the conclusions are available in the paper or the Supplementary Materials. Raw data and computer codes used for data analysis are available from the corresponding author.

## Experimental Model and Subject Details

### Cell lines and tissue culture

HEK293T, Plat-E, and mouse embryonic fibroblasts were cultured using DMEM (Genesee Scientific 25-500) + 10% FBS (Genesee Scientific 25-514H) and incubated at 37C with 5% CO2. Plat-E cells were selected using 10 µg/mL blastocidin and 1 µg/mL puromycin every three passages. HEK293T, Plate-E, and mouse embryonic fibroblasts were detached from flasks using Trypsin (Genesee Scientific 25-510) diluted in PBS. For seeding and passaging, cells were counted using a hemocytometer. Cells were periodically tested for mycoplasma.

### iPSC cell culture

IPS11 cells were cultured using mTeSR™ Plus (STEMCELL Technologies, 100-1130) with Geltrex™ (ThermoFisher Scientific, A1413302) coating and incubated at 37C with 5% CO2. For passaging, iPS11 cells were dissociated in clumps using ReLeSR™ (STEMCELL Technologies, 100-0484) according to the manufacturer’s instructions. For experiments, iPS11 cells were dissociated into single cells using Gentle Cell Dissociation Reagent (STEMCELL Technologies, 100-1077), counted using a hemocytometer, and replated in mTeSR™ Plus with ROCK inhibitor (Millipore Sigma, Y0503-5MG) and Penicillin-Streptomycin (ThermoFisher Scientific, 10-378-016).

### MEF dissection and isolation

C57BL/6 mice were mated with mice bearing the Hb9::GFP reporter. Mouse embryonic fibroblasts were isolated at E12.5-E14.5 under a dissection microscope. Embryos were sorted into non-transgenic and Hb9::GFP+ by using a blue laser to illuminate the spinal cord to identify the presence of Hb9::GFP+ cells. After removing the head and internal organs to avoid contaminating neurons and other cells, razors were used to break up the tissue for 5 minutes in the presence of 0.25% Trypsin-EDTA. Up to two embryos were processed at the same time. The preliminary suspension was neutralized, resuspended, and triturated with 0.25% Trypsin-EDTA. The suspension was neutralized and resuspended again before filtering through a 40 µm cell strainer. MEF cultures were expanded on 10 cm dishes coated with 0.1% gelatin, using one 10 cm dish per embryo. MEFs were expanded until ∼80% confluent, passaged, and expanded for at least 3-4 days. Passage 1 MEFs were tested for mycoplasma, cryopreserved in 90% FBS and 10% DMSO, and stored in liquid nitrogen.

## Method Details

### Plasmid Cloning

Plasmids were cloned using standard cloning protocols for Gibson assembly and Golden Gate assembly. Promoters, target genes, and polyA signals were inserted into part position vectors (pPVs) via Gibson assembly. Expression plasmids were assembled via BsaI (New England Biolabs) Golden Gate Assembly using pPVs or Q5 polymerase-amplified and gel-purified DNA fragments. The lentivirus or retrovirus plasmids were assembled with PaqCI Golden Gate Assembly using expression plasmids and viral backbone plasmids.

### DIAL promoter cloning

The DIAL promoter was constructed by inserting a spacer sequence between loxP or VloxP sites. The spacer is a tandem sequence of two putatively neutral regions of an expression plasmid, joined by restriction digest and Gibson assembly. The loxP and VloxP sequences were taken from respective recombinase reporters (gifts from the Wong Lab at Boston University, BW338 and BW273). Tessellated ZF43/37 hybrid binding sites were taken from ZF reporter Addgene plasmids #138934. The DIAL promoter was assembled through Golden Gate or Gibson assembly, using PCR products of spacer sequence and ZF binding sites with complementary overhangs. Additional recombinase binding sites were inserted using Gibson assembly with PCR overhangs. The assembled DIAL promoters were inserted into pPVs to produce downstream gene constructs through Golden Gate assembly.

### TET-DIAL promoter cloning

The TET-DIAL promoter was constructed by inserting a spacer sequence between loxP sites. The spacer is a putatively neutral region from an expression plasmid. The loxP sequences were taken from recombinase reporter (gift from the Wong Lab at Boston University, BW338). TetO binding sites were taken from TRE-dCas9-VPR Addgene plasmid #63800. The TET-DIAL promoter was assemblyed through BsaI Golden Gate assembly, using PCR products of the spacer, loxP recombinase sites, TetO binding sites and expression vector with complementary overhands. Additional loxP sites were inserted via annealed oligos with complementary overhangs via BsaI Golden Gate assembly. All TET-DIAL promoters were cloned into pSHIPs for expression in transfection of HEK293T cells.

### Recombinase and Zinc Finger Activator Plasmids

Unless otherwise specified, in transient transfections, recombinase was expressed as: Cre from CAG-Cre-RbG (Addgene plasmid #89573), VCre from CAG-VCre-RbG (Addgene plasmid #89575), GIB-Cre N-terminal (Addgene plasmid #108723), and GIB-Cre C-terminal (Addgene plasmid #108724) (gifts from the Wong Lab at Boston University). Unless otherwise specified, zinc finger activators (ZFa) were expressed from EF1a-VP16-ZF37-2A-mCherry-BGH, EF1a-VP16-ZF43-2A-TagBFP-BGH, CMV-VP64-ZF37-BGH (Addgene plasmid #138834), CMV-VP64-ZF43-BGH (Addgene plasmid #138835), and CMV-VPR-ZF37-BGH (Addgene plasmid #138839). PCR fragments of Addgene plasmids #138759 and #138730 were used to clone target gene pPVs for VP16-ZF37 and VP16-ZF43 through Gibson assembly. The 2A-tagged ZFs were subsequently inserted into expression vectors with EF1a and BGH polyA signal via Golden Gate assembly with BsaI (NEB Stable).

### HEK293T DNA Transient Transfection

39,000-42,000 cells were plated per well in a 96-well plate 24 hours before transfection. Cells were transfected on day 0 with 112.5 ng/plasmid/well delivered via a 4:1 ratio of ug PEI to ug of DNA mixed in KnockOut DMEM (Fisher Scientific, 10-829-018). Unless specified, each well was transfected with 112 ng of all plasmids except the recombinase plasmid, which was transfected with 11 ng per well. All transfection experiments included a co-transfection marker EF1a-iRFP670-BGH (112 ng). Also, all transfection experiments had the same total DNA and PEI per condition within the experiment by using an empty vector plasmid to keep the total ug of DNA per well consistent. At 1-day post-transfection (dpt), media was aspirated and replaced with DMEM + 10% FBS. As specified in figure captions, in some experiments, this replacement media at 1 dpt contained 1x DOX (1 ug/mL, from 1000X stock of 1 mg/mL stored at -20C or 4C) and appropriately titrated down conditions or 1x gibberellin (G3-AM, Sigma Aldrich, from 1000X stock of 10 mM dissolved in ethanol and stored at -20 C). For applicable experiments, modRNA was also transfected at 1 dpt. Cells were flowed and imaged at 3 dpt in technical triplicates. Bioreplicates represent separate transfection experiments. All experiments were conducted with biological replicates (n=3 or more). To detach cells from wells in preparation for flow cytometry, 1:1 Trypsin/1xPBS mixture was added to each well. After 8 minutes, DMEM + 10% FBS was added on top before centrifuging down the plates at 1000 rpm for 10 minutes. Media was aspirated and cells were resuspended in 1x PBS to transfer to a round bottom plate for flow cytometry. Unless specified, transfection refers to transient transfection of plasmid DNA. DNA amounts are detailed in Table 1.

### iPSC Transient Transfection Experiments

15,000 iPS11 cells were plated per well in a 96-well plate 48 hours before transfection. 24 hours before transfection, media containing ROCK inhibitor was removed and replaced with mTeSR™ Plus with Penicillin-Streptomycin. On the day of transfection, the media was changed to Opti-MEM™ (ThermoFisher Scientific, 31985062) and transfection mixes were prepared with FUGENE® HD (FuGENE, HD-1000) according to the manufacturer’s instructions (ratio of 3 uL reagent to 1 ug DNA). Each well was transfected with 100 ng plasmid reporter, 12.5 ng zinc finger plasmid, 25 ng Cre plasmid, and 25 ng transfection marker plasmid. At 4 hours after transfection, mTeSR™ Plus with Penicillin-Streptomycin was added. At 24 hours after transfection, the media was changed to mTeSR™ Plus with Penicillin-Streptomycin and if specified DOX (1 µg/mL). Cells were flowed at 2 DPT and imaged at both 1 DPT and 2 DPT in technical triplicates. Bioreplicates represent separate transfection experiments. All experiments were conducted with biological replicates (n=3). To flow, cells were detached from wells using Gentle Cell Dissocation Reagent (STEMCELL Technologies, 100-1077) for 8 minutes and centrifuged at 500g for 5 minutes. Media was aspirated and cells were resuspended in 1x PBS and transferred to a round bottom plate for flow cytometry. Unless specified, transfection refers to transient transfection of plasmid DNA. DNA amounts are detailed in Table 1.

### PCR of Cell Lysis

DNA from transfected HEK293Ts was obtained with Cell Lysis Buffer (10X) (Cell Signaling Technology 9803S). Cells were detached from wells using a 1:1 Trypsin/1x PBS mixture for 8 minutes. DMEM + 10% FBS was added on top and plates were centrifuged down at 1000 rpm for 10 minutes. Media was aspirated and cells were resuspended in 50 uL Cell Lysis Buffer (10X) and 0.5 uL of Proteinase K (New England Biolabs P8107S) per 96-well. Cells were incubated for 45 minutes at 85 C then microfuged. 1 uL of the supernatant was used for PCR using Apex Taq RED Master Mix, 2X (Genesee Scientific 42-138B). PCR products were run on a 2% agarose gel at 110V for 45-60 minutes.

### Retrovirus production in Plat-Es

Plat-E cells were seeded at 850k per 6-well onto plates coated with 0.1% gelatin. The next day, Plat-Es were transfected with 1.8 µg of DNA per well using a 4:1 ratio of µg PEI:µg DNA. The next day, the media was replaced with 1.25 mL fresh 25 mM HEPES buffered DMEM with 10% FBS. The next day, viral supernatant was collected and filtered through a 0.45 µm filter. 1.25 mL fresh media was again added to the Plat-E cells. Filtered viral supernatant was then used for transduction of MEFs. Viral supernatant collection, filtration, and transduction was repeated for a second day. One day after the second viral transduction was considered one day post infection (dpi).

### Lentiviral production in HEK293Ts

HEK293T cells were seeded at 1 million per 6-well onto plates coated with 0.1% gelatin. The next day, each well of 293Ts was transfected with 1.02 µg packaging plasmid (psPax2), 2.05 µg of envelope plasmid (pMD2.G), and 1.02 µg of transfer plasmid using a 4:1 ratio of ug PEI:μg DNA. 6-8 hours later, the media was replaced with 1.25 mL of fresh 25 mM HEPES buffered DMEM with 10% FBS. 24 and 48 hours later viral supernatant was collected and stored at 4°C, replacing with fresh media after the first collection. After the second collection, the viral supernatant was filtered with a 0.45 μM filter and was mixed with 1 volume Lenti-X concentrator to 3 volumes viral supernatant and incubated at 4°C overnight. The virus was pelleted by centrifugation at 1,500 x g for 45 minutes at 4°C. The supernatant was removed, and the pellets were resuspended in media to a final volume of 33 μL per 6-well of virus. Volumes and cell numbers were scaled up proportionally for a 10-cm plate. Concentrated virus was resuspended to a final volume of 1 mL per 10-cm plate of virus. The virus was used for transduction of MEFs on the same day.

### Viral transduction of mouse embryonic fibroblasts (MEFs)

MEFs were seeded one day prior to viral transduction onto plates coated with 0.1% gelatin at 10k per 96-well. MEFs were transduced two days in a row with 11 μL of each PlatE retrovirus per 96-well. On the second day MEFs were also transduced with 3 µL per 96-well of concentrated lentivirus from a 6-well plate, or 5 uL per 96-well of concentrated lentivirus from a 10-cm plate. Fresh media was included to reach a final volume of 100 μL per 96-well, and 5 ug/mL polybrene was added to increase transduction efficiency. One the second day of transduction the plates were also centrifuged at 1500 x g for 90 minutes at 32°C to further increase lentiviral transduction efficiency. At 1 dpi the transduction media was replaced with fresh DMEM + 10% FBS. If specified in figure caption, DOX (1 µg/mL) was also added at 1 dpi. At 3 dpi, the cells were dissociated using trypsin and analyzed via flow cytometry.

### In vitro transcription and transfection of modRNA

The plasmid templates used for Cre and TagBFP modRNA synthesis harbors the 5’-UTR of human β-globin, a Kozak sequence, the coding sequence for iCre, and the 3’-UTR of human β-globin. The linear template for *in vitro* transcription (IVT) was generated via PCR using Q5 DNA Polymerase (New England Biolabs) with the forward primer (5’-AGCTATAATACGACTCACTATAAGctcctgggcaacgtgctg-3’) encoding the T7 promoter (upper-case bases) and binding the 5’ UTR (lower-case bases) and the reverse primer (5’-poly(T)_116_-GCAATGAAAATAAATGTTTTTTATTAGGCAGAAT-3’) encoding the poly(A) tail and binding the 3’-UTR. The PCR product was isolated on a 1% agarose gel, excised, and purified using the Monarch PCR and DNA Cleanup Kit (New England Biolabs). 200 ng of purified product was used in a 20 µL IVT reaction using the HiScribe T7 High Yield RNA Synthesis Kit (New England Biolabs), fully substituting UTP with N1-methylpseudouridine-5’-phosphate (TriLink Biotechnologies) and co-transcriptionally capping with CleanCap Reagent AG (TriLink Biotechnologies). IVT reactions were incubated at 37 °C for 4 hr, at which point reactions were diluted to 50 µL, treated with 2 µL DNase I, and incubated at 37 °C for 30 min to degrade the IVT PCR template DNA. Synthesized modRNA was column purified and eluted with 60 µL water using the 50 µg Monarch RNA Cleanup Kit (New England Biolabs). A small sample was nanodropped and ran on a native denaturing gel to determine RNA concentration and verify full-length product. The modRNA was dispensed in single-use aliquots and stored at –80 °C. For Cre modRNA and TagBFP modRNA transfection experiments, each 96-well was transfected with 200 ng of modRNA total (100 ng of each) using 0.4 µL Lipofectamine MessengerMAX (Thermo Fisher Scientific) according to manufacturer’s instructions.

### Viral induction of HEK293T Cells, Cell Line Development, and modRNA Transfection

HEK293T cells were seeded on the day of viral transduction in suspension at 20,000 cells per 96-well. Each 96-well was transduced with 3 µL per concentrated lentivirus from a 6-well plate. Fresh DMEM with 10% FBS was included to reach a final volume of 100 μL per 96-well, and 5 ug/mL polybrene was added to increase transduction efficiency. The co-infected lentiviruses were: lentivirus containing ZFa cassette EF1a-VP16-ZF37-2A-mCherry-BGH and lentivirus containing with 203 bp spacer DIAL promoter with mGL-WPRE, with and without divergent EF1a-iRFP670-BGH. At 3 dpt, cells were passaged into a 6-well plate. At 10 dpt, cells positive for mCherry and mGL were sorted using a Sony MA900. Cells were cultured in a 24-well plate until confluent. From this polyclonal population, 20,000 cells then plated into six 96-wells. One condition in triplicate was transfected with 100 ng of TagBFP modRNA and 100 ng of Cre modRNA. The other condition was not transfected. At 3 dpt, cells were analyzed via flow cytometry. A subset of one-fifth of the cells were re-plated. These passaged cells were then analyzed via flow cytometry at 6 dpt, with a subset of one-fifth of the cells replated again. These passeges cells were then analyzed via flow cytometry at 9 dpt. Flow data was gated for both mGL and mCherry for analysis.

### Imaging

Images were taken on Keyence™ All-in-one fluorescence microscope BZ-X800.

### Flow cytometry data analysis

Live cells were gated based of FSC-A and SSC-A using FlowJo. Subsequently, single cells were gated via SSC-H and SSC-A. Single cells were then exported all plots and statistics were generated from pandas, matplotlib, and seaborn packages in Python. For transient transfection of all HEK293T cells, cells were additionally gated by co-transfection marker iRFP670 as shown in Figure S2, at iRFP670-A>10000 or at iRFP670-A>20000 always consistent within each plot and experiment. Additional gating strategy for virally-delivered cassettes or modRNA are described in the figure captions. For transient transfection of all iPSCs, cells were additionally gated by co-transfection marker TagBFP with a similar method as Figure S2, at TagBFP-A>4000 for all experiments. Further gating based on ZFa co-expressed fluroscent proteins or output positive cells in transient transfection is specified in figure captions.

### Mathematical modeling of DIAL activity for varying levels of ZFa

Dose-dependent reporter gene activation via ZFa titration was modeled using the Hill equation.

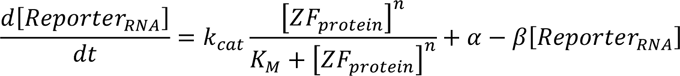

The relationship between reporter gene transcription and translation was simplified to obtain

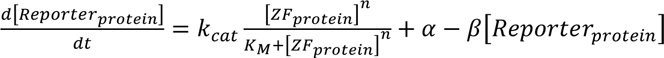

where *k*_*cat*_ is the translation rate constant; α is the leaky expression from TF-independent transcription; and β is the degradation constant.

Assuming steady state of cellular processes, the reporter protein levels are expressed as

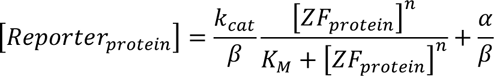

Experimental data from the ZFa was divided into conditions (e.g in the presence or absence of Cre) to generate separate parameters that captures the different gene regulation across pre- and post-excision constructs. Parameters *k*_*cat*_ and α consolidated to simplify the steady state equation as follows:

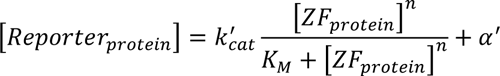

The leakiness parameter α^′^ was set to the lowest value of output mGL in each condition corresponding to no ZFa added. Assuming no Hill cooperativity for ZF binding (*n* = 1), the experimental data was used as [*ZF_protein_*] and [*Reporter_protein_*] to calculate parameters 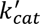 and *K_M_* from the equation.

To minimize technical variance, the ZFa titration data was normalized by the 0.125xZFa condition (14 ng per well) within each bioreplicate. Potentially due to resource burden at high DNA concentration of ZFa, expression of mGL decreased at the highest levels of input mCherry (ZF37). These datapoints were removed before generating parameters to prevent the predicted parameters from reflecting resource burden which is not incorporated in these model equations. The data was transformed by the leaky mCherry or TagBFP expression in conditions with no ZF to normalize 0x ZFa conditions to no input ZF expression levels.

To fit the model, the data was bootstrapped across each condition (e.g in the presence or absence of Cre). The level of protein expression from the DIAL promoter was used to calculate parameters and generate a corresponding predicted titration curve for each bootstrapped sample. The 95% confidence interval of the titration curve was generated by selecting the 2.5^th^ and 97.5^th^ percentile output mGL values for each input mCherry or TagBFP value.

## Figures

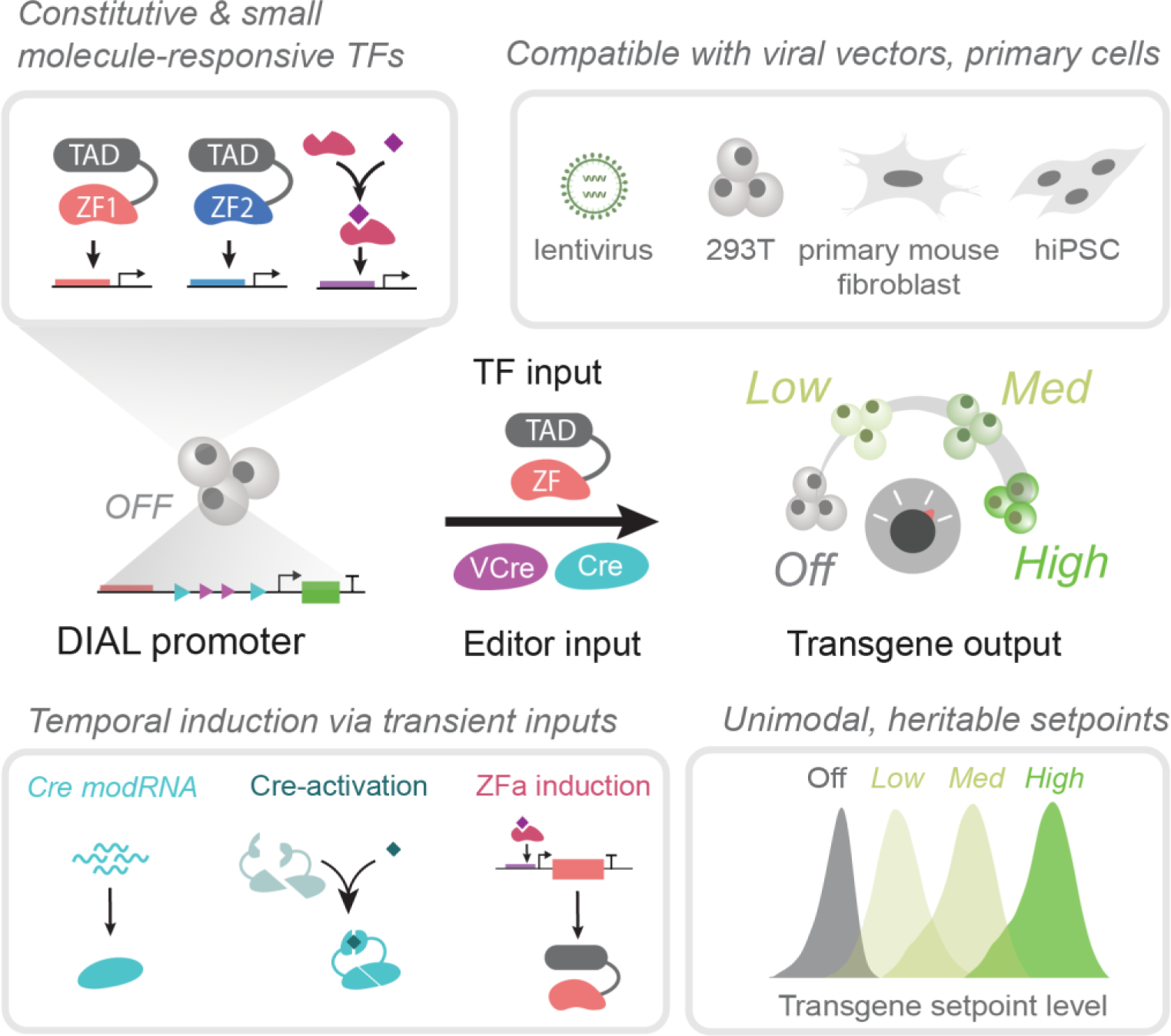

## Overview

DIAL offers an extensible framework for designing synthetic promoters that generate heritable setpoints of gene expression and perform across a range of cell types and delivery systems.

