## Supplement for "Programmable promoter editing for precise control of transgene expression"

### Supplementary Figures

**Figure S1. DIAL promoter changes the setpoint by reducing the distance between the binding sites and minimal promoter.**

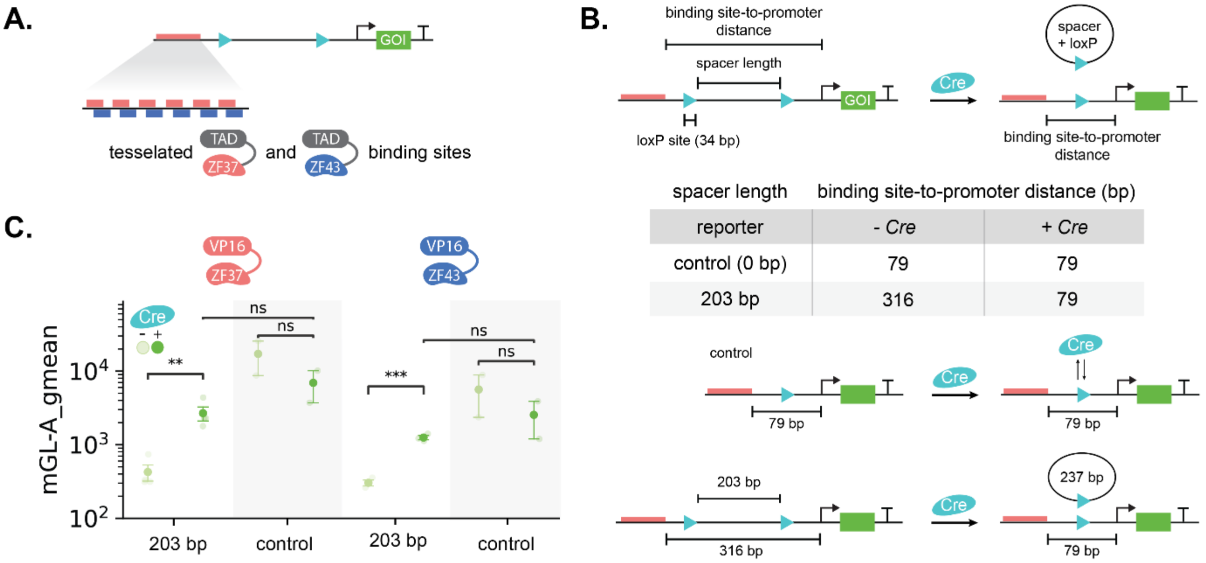


(A) Schematic of tessellated hybrid binding sites for ZF37 and ZF43 placed upstream of the floxed spacer to build the DIAL promoter. The DIAL system is composed of the DIAL promoter, recombinase, and the cognate ZFa that induces expression of the gene of interest.

(B) Schematic of the pre- and post-excision states of the DIAL promoter. Spacer length is defined as the distance between the loxP recognition sites. The binding site-to-promoter distance is defined as the total distance between the last binding site and the minimal promoter. Cre excises a loxP site (34 bp) along with the spacer. The control statically encodes the post-excision promoter state. In the control, the binding site-to-promoter distance is 79 bp. While Cre may bind to the remaining loxP site on the post-excision constructs, editing does not occur.

(C) Geometric mean fluorescence intensity (MFI) of mGL expressed from the 203 bp DIAL promoter or control transfected on plasmids into HEK293T cells. In the presence of Cre for both ZFas, the output of the DIAL promoter converges to the output level of the control. Addition of Cre to the control promoter slightly reduces output expression.

All units for output MFI are arbitrary units (a.u.), and fold change is unitless. Large markers represent the mean of biological replicates with span indicating standard error (n=3).

**Figure S2. Gating strategy for isolating transfected cells based on the co-transfection marker.**

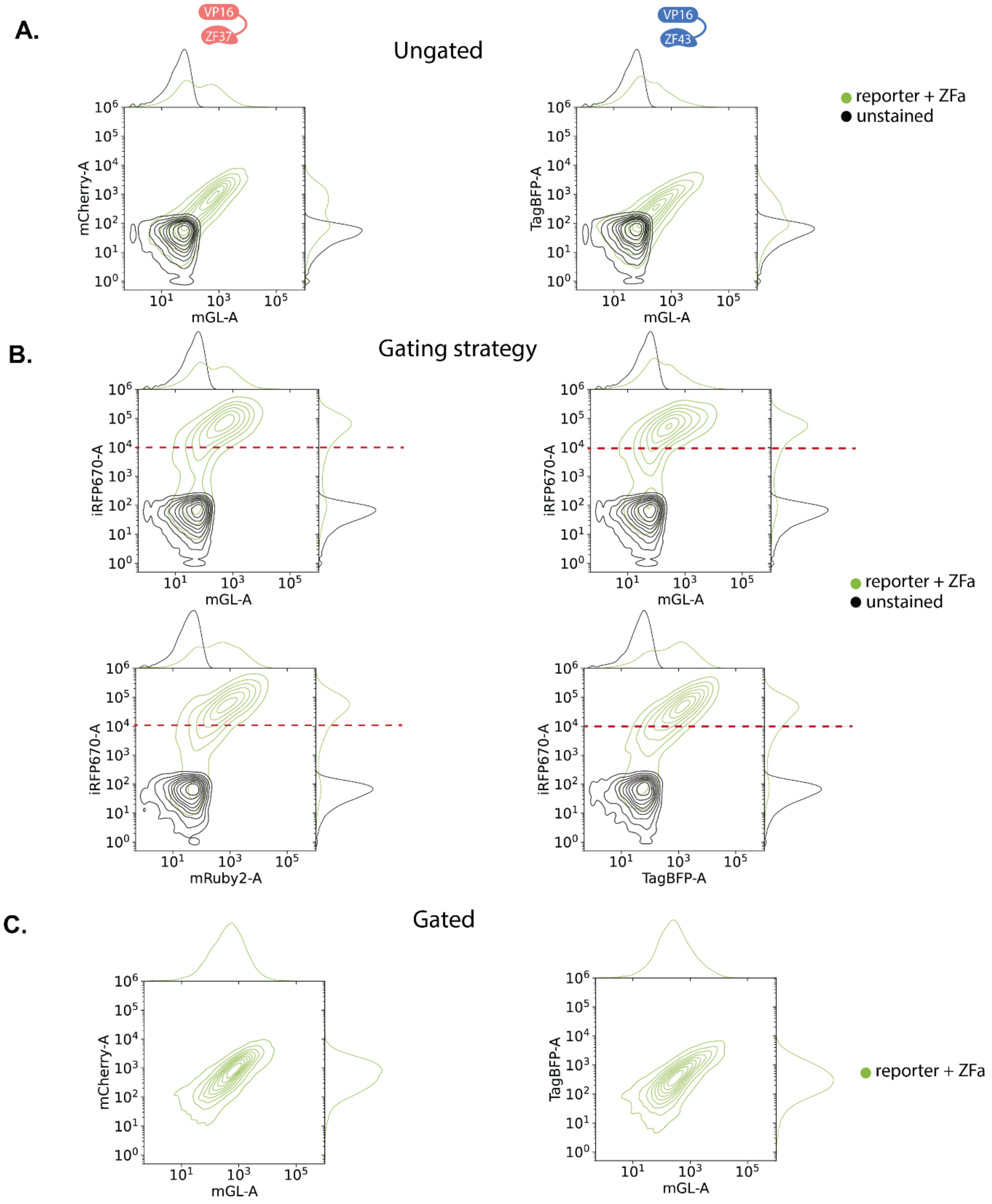


(A-C) Histograms and joint contour plots of mGL expressed from the 203 bp DIAL promoter, co-transfected ZFa (co-expressed with mCherry or TagBFP), and separate transfection marker iRFP670 on plasmids transfected into HEK293Ts. With ZFa conditions correspond to Fig 1C and S1C. Histograms and joint contour plots represent single-cell distributions sampled across bioreplicates (n=3). Light green: ZFa (VP16-ZF37 or VP16-ZF43) with the DIAL promoter. Black: Untransfected HEK293T cells.

(A) Populations gated for live, single cells. The ZFa protein expression levels are proxied by the fluorescence intensity of mCherry (co-expressed with VP16-ZF37) or TagBFP (co-expressed with VP16-ZF43). Output level of the DIAL promoter is represented by mGL fluorescence intensity.

(B) Gating strategy of live, single cells based on iRFP670 marker. Gate shown as dashed lines at the threshold of the co-transfection marker iRFP670 > 10,000.

(C) Live single cells gated to be positive for iRFP670. When controlling for transfection, the distribution of both ZFa (proxied by TagBFP or mCherry) and output reporter gene (mGL) is unimodal.

**Figure S3. No expression from the DIAL promoter in the absence of ZFa. ZFa unaffected by Cre.**

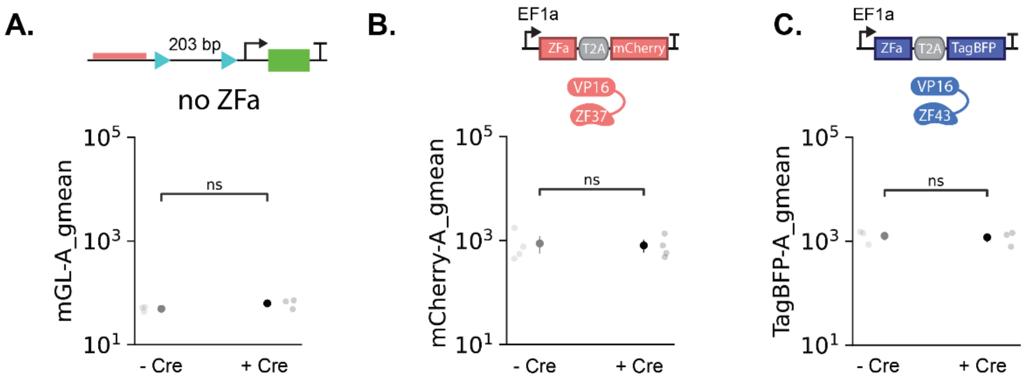


(A) Geometric mean fluorescence intensity (MFI) of mGL expressed from the 203 bp spacer DIAL promoter transfected on plasmids into HEK293T cells without ZFa, and with or without Cre. In the absence of ZFa, the output of the DIAL promoter is unaffected by the presence of Cre.

(B) Geometric MFI of mCherry coexpressed with ZFa (VP16-ZF37) transfected on plasmids into HEK293T cells along with the 203 bp spacer DIAL promoter and with or without Cre. The presence of Cre does not affect the ZFa levels.

(C) Geometric MFI of TagBFP coexpressed with ZFa (VP16-ZF43) transfected on plasmids into HEK293T cells along with the 203 bp spacer DIAL promoter and with or without Cre. The presence of Cre does not affect the ZFa levels.

All units for output MFI are arbitrary units (a.u.). Large markers represent the mean of biological replicates with span indicating standard error. Shaded points represent individual bioreplicates (n=3 for A and C; n=4 for B).

**Figure S4. Cre-mediated DNA editing changes the state of the DIAL promoter.**

**A.**

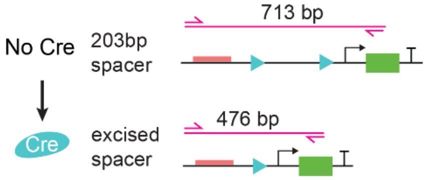


**C.**

| spacer length | PCR product length | |
|---|---|---|
| reporter | *- Cre* | *+ Cre* |
| control (0 bp) | 476 | 476 |
| 27 bp | 537 | 476 |
| 79 bp | 589 | 476 |
| 155 bp | 665 | 476 |
| 203 bp | 713 | 476 |
| 263 bp | 773 | 476 |

**B.**

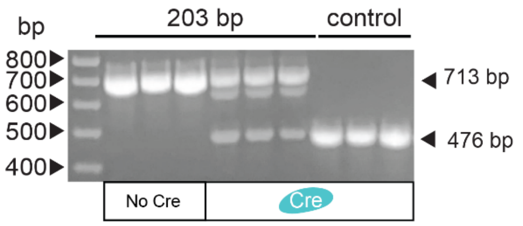


**D.**

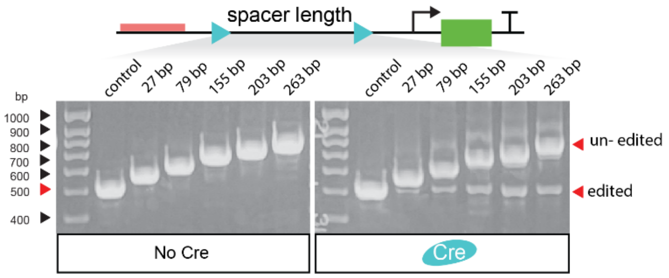


(A) Schematic of PCR used to visualize changes to the state of the DIAL promoter. The primers bind to sequences upstream of the ZF binding sites or the mGL reporter gene. In the presence of Cre, the spacer excision shortens the distance between binding sites and the minimal promoter, resulting in a shorter PCR product.

(B) Image of gel electrophoresis of PCR products using the primers shown in A. PCR was performed on cell lysis from HEK293T cells transfected with combinations of plasmids for Cre, the 203 bp spacer or control DIAL promoter, and VP16-ZF37. The first three lanes after the ladder are PCR products of conditions with 203 bp spacer DIAL promoter and VP16-ZF37. The six following lanes are PCR products with control or 203 bp spacer DIAL promoter, VP16-ZF37, and Cre. Samples are technical replicates (n=3). The condition with 203 bp spacer DIAL promoter and Cre (middle 3 lanes) shows a shorter band at 476 bp, the expected length of the PCR product from the post-excision promoter.

(C) Table of expected sizes of PCR product from control and different spacer length DIAL promoters, with and without Cre. According to our promoter design, DIAL promoters of all spacer lengths should have the same size PCR product for its post-excision state.

(D) Image of gel electrophoresis of PCR products using the primers shown in (A). PCR was performed on cell lysis from HEK293T cells transfected with combinations of plasmids for Cre, control or different spacer length DIAL promoters, and VP16-ZF37. The left gel shows PCR products of conditions with control or different spacer length DIAL promoters regulating mGL with VP16-ZF37 and without Cre. The right gel shows PCR products of conditions with control or different spacer length DIAL promoters regulating mGL with both VP16-ZF37 and Cre. The PCR products in the right gel include a band around 476 bp, suggesting the emergence of a post-excision promoter state in the presence of Cre.

**Figure S5. DIAL promoter system is tuned by spacer length and ZFa identity.**

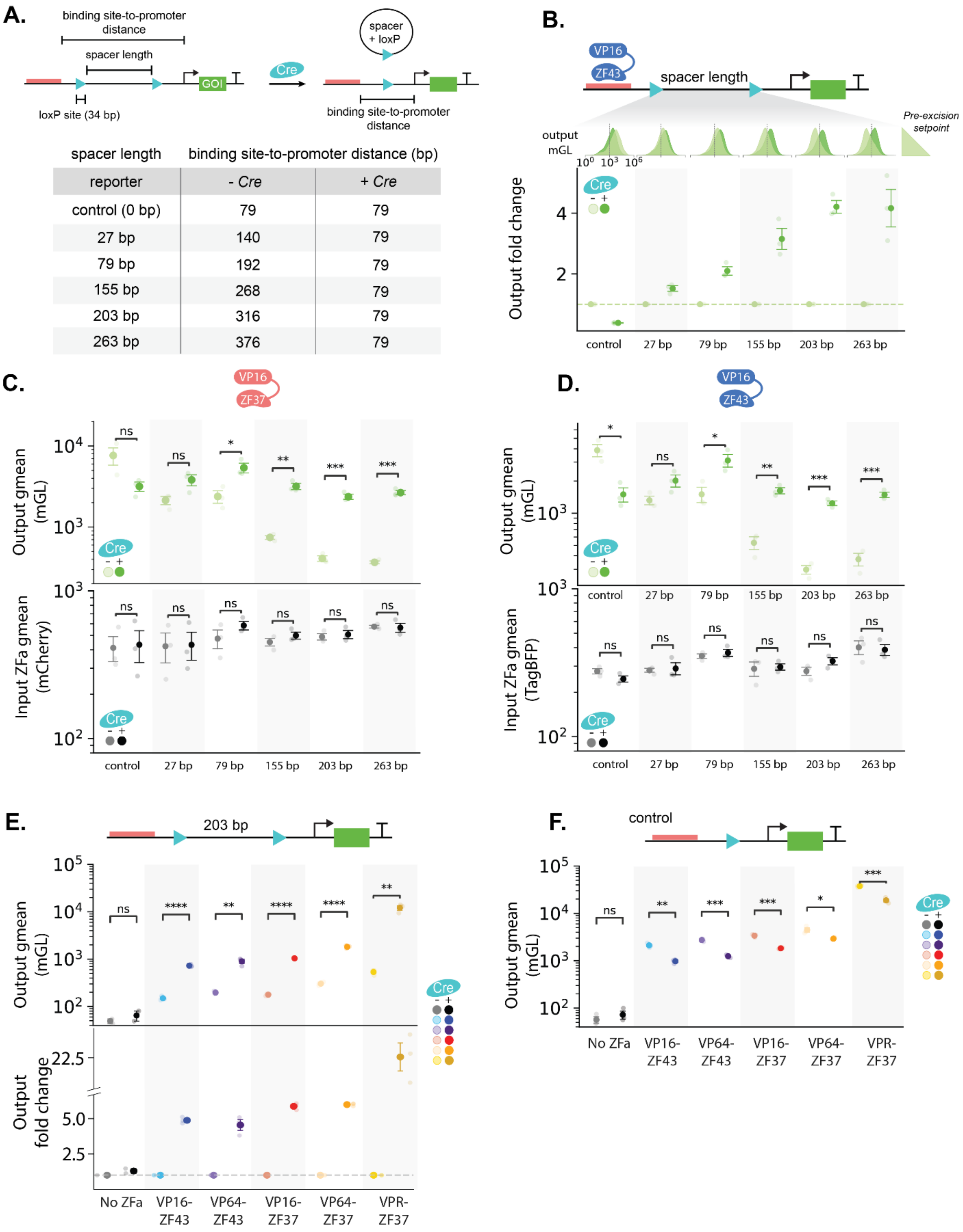

A. Schematic of the pre- and post-excision molecular states of the DIAL promoter. The table summarizes the binding site-to-promoter distances for DIAL promoters with different spacer lengths, with and without Cre.

B. Fold change of the output reporter expressed from DIAL promoters with different spacer lengths transfected with ZFa (VP16-ZF43) on plasmids into HEK293T cells, in the presence or absence of Cre. Fold change is the output mGL geometric mean fluorescence intensity (MFI) normalized to the condition without Cre within each spacer length. Histograms show decreasing pre-excision expression for increasing spacer length, which generates the larger fold change upon addition of Cre.

C. Geometric MFI of output mGL expressed from DIAL promoters and mCherry (coexpressed with ZFa VP16-ZF37). Pre-excision expression decreases with increasing spacer length. In the presence of Cre, the output of the DIAL promoter converges to the output level of the control. ZFa levels (proxied by mCherry geometric MFI) are not affected by adding Cre nor increasing spacer length.

D. Geometric MFI of output mGL expressed from DIAL promoters and TagBFP (coexpressed with ZFa VP16-ZF43). Pre-excision expression decreases with increasing spacer length. In the presence of Cre, the output of the DIAL promoter converges to the output level of the control. ZFa levels (proxied by TagBFP geometric MFI) are not affected by adding Cre nor increasing spacer length.

E. Geometric MFI of output mGL expressed from DIAL promoters transfected on plasmids into HEK293T cells with different zinc finger activators bearing different transactivation domains (ZF-TADs or ZFas), with or without Cre. Fold change is the output mGL geometric MFI normalized to the condition without Cre within each ZF-TAD. Expression setpoints and fold changes increase slightly with stronger ZF-TAD.

F. Geometric MFI of output mGL expressed from DIAL promoters transfected on plasmids into HEK293T cells with different zinc finger activators bearing different transactivation domains (ZF-TADs), and with or without Cre. Expression setpoints increase slightly with stronger ZF-TAD. Presence of Cre decreases expression.

All units for output MFI are arbitrary units (a.u.), and fold change is unitless. Large markers represent the mean of biological replicates with span indicating standard error. Histograms represent single-cell distributions sampled across bioreplicates (n=3). Statistical significance was calculated with Students t-Test with ns $p>0.05$; *$p<0.05$; **$p<0.01$; *** $p<0.001$.

**Figure S6. The nested DIAL promoter generates three promoter states and four states of expression.**

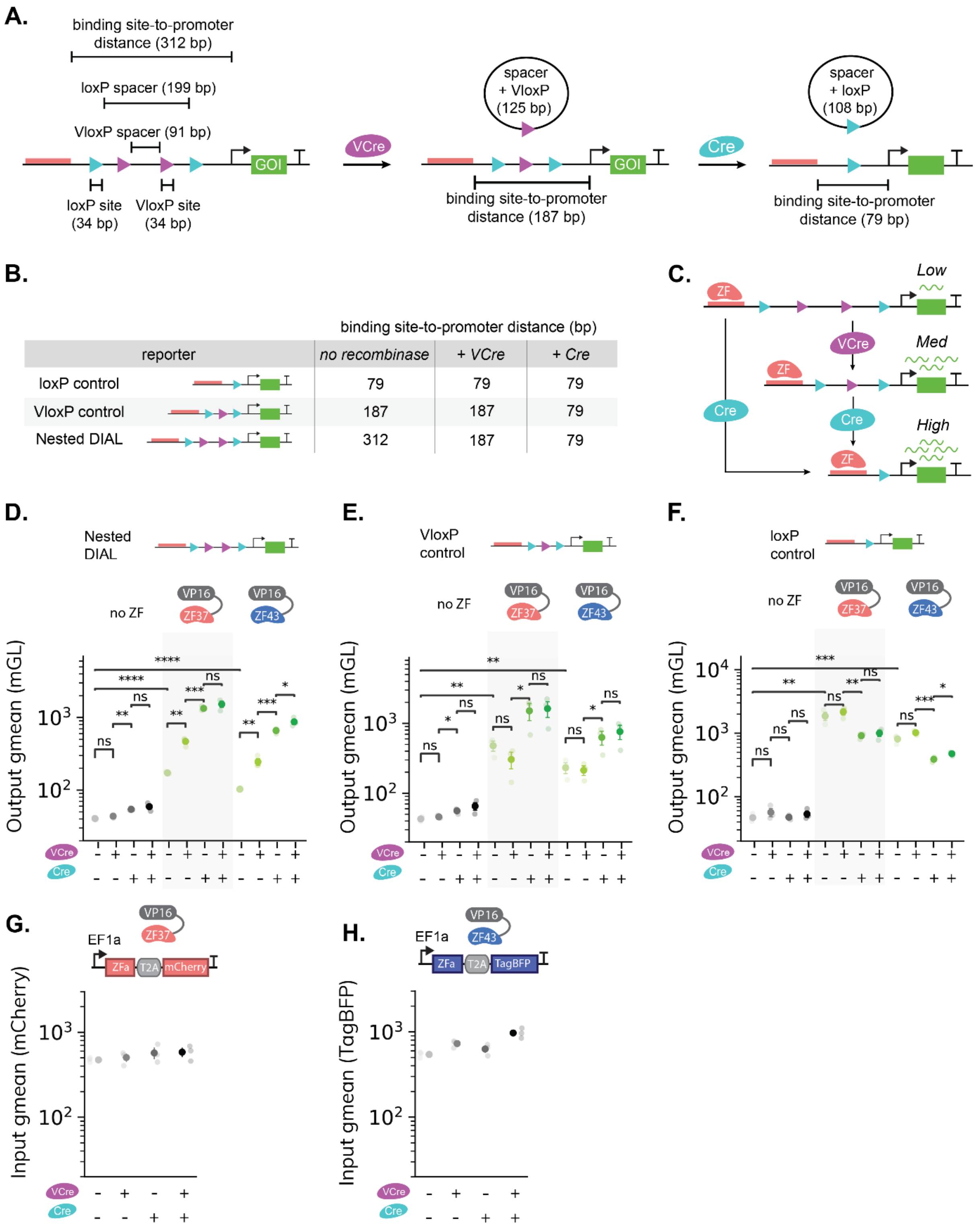

(A) Schematic of the pre- and post-excision states of the nested DIAL promoter. VloxP and loxP spacers are defined as the distance between the VloxP and loxP recognition sites, respectively. The binding site-to-promoter distance is defined as the total distance between the last binding site and the minimal promoter. VCre and Cre excise a VloxP and loxP site along with the spacer, respectively. The nested DIAL promoter has three possible promoter states.

(B) The table summarizes the binding site-to-promoter distances for the loxP control, VloxP control, and nested DIAL promoters, with or without VCre and Cre. The VloxP and loxP control statically encodes the post-excision promoter state after adding VCre and Cre, respectively. Adding VCre to VloxP control and adding Cre to loxP control does not edit the promoter, not changing the binding site-to-promoter distance.

(C) Schematic of nested spacer achieving three different expression setpoints (excluding "off" state without ZFa). In the presence of ZFa, he nested spacer can go through all three promoter states and corresponding setpoints by sequentially adding VCre then Cre, or skip directly to the "high" setpoint by adding Cre.

(D,E,F) Geometric MFI of mGL expressed from the nested, VloxP control, and loxP control DIAL promoter transfected on plasmids into HEK293T cells with and without ZFa (VP16-ZF37 or VP16-ZF43) or Cre. In the absence of ZF, the outputs of the DIAL promoter are negligible and not substantially different.

(D) In the presence of ZFa and VCre, the output of the nested DIAL promoter increases and converges to the output level of the VloxP control DIAL promoter shown in E. In the presence of ZFa and Cre, the output of the nested DIAL promoter increases converges to the output level of the loxP control DIAL promoter shown in F.

(E) In the presence of ZFa, adding VCre slightly reduces the output of the VloxP control DIAL promoter. Adding Cre increases the output, converging to the output level of the loxP control shown in F.

(F) In the presence of ZFa, adding Cre slightly reduces the output of the loxP control DIAL promoter, consistent with previous data. This decrease is not observed with the addition of VCre.

(G) Geometric mean fluorescence intensity (MFI) of mCherry co-expressed with ZFa (VP16-ZF37) transfected on plasmids into HEK293T cells along with the nested DIAL promoter with or without Cre. The presence of recombinase does not substantially affect ZFa levels.

(H) Geometric mean fluorescence intensity (MFI) of TagBFP co-expressed with ZFa (VP16-ZF43) transfected on plasmids into HEK293T cells along with the nested DIAL promoter with or without Cre. The presence of recombinase does not substantially affect ZFa levels.

All units for output MFI are arbitrary units (a.u.), and fold change is unitless. Large markers represent the mean of biological replicates with span indicating standard error (n=3). Statistical significance was calculated with Students t-Test with ns $p>0.05$; *$p<0.05$; **$p<0.01$; *** $p<0.001$.

**Figure S7. DIAL robustly generates expression setpoints with VP16-ZF43.**

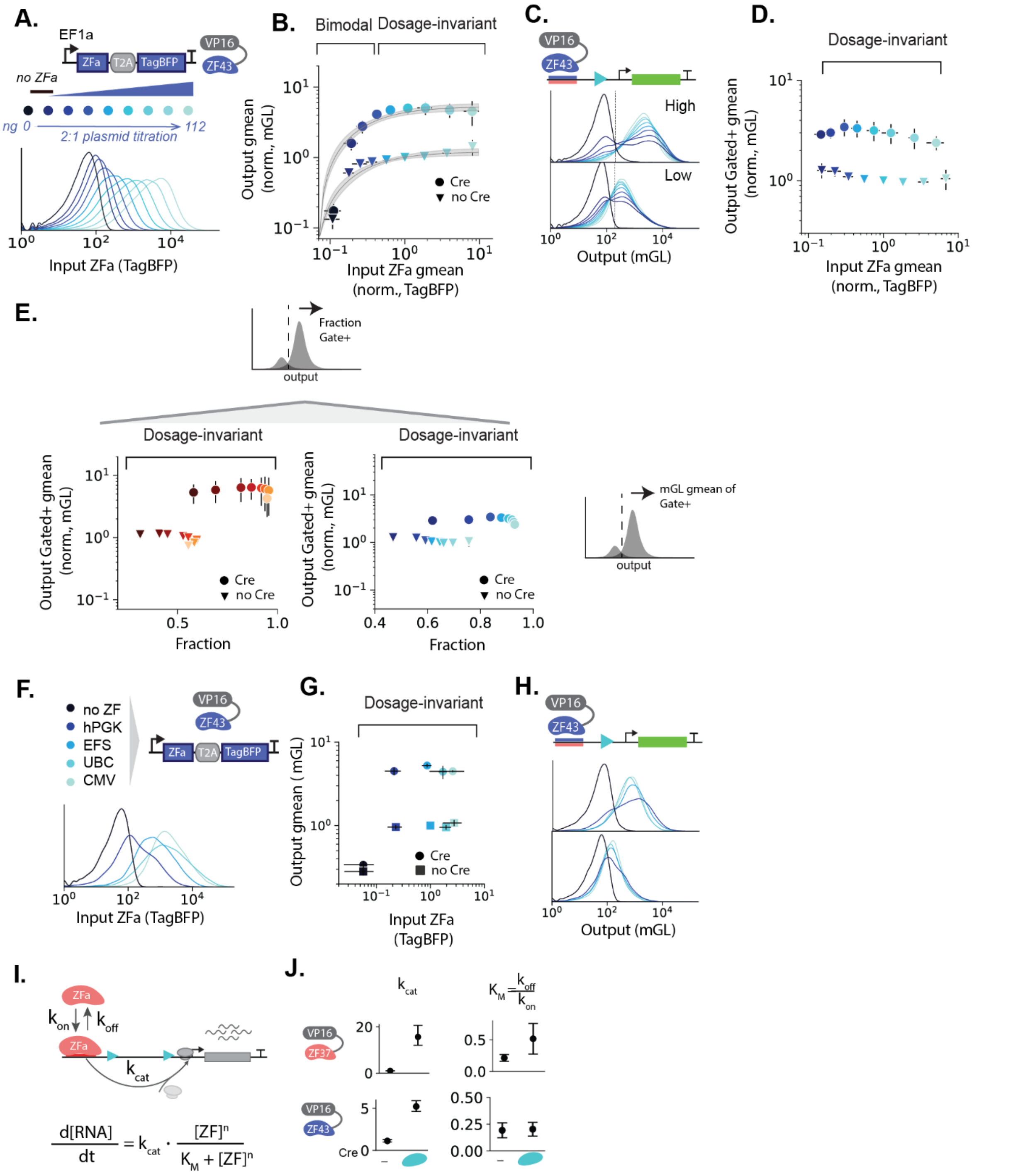


(A) Single-cell distributions of TagBFP coexpressed with ZFa (VP16-ZF43) transfected on plasmids into HEK293T cells with 203 bp DIAL promoter. Conditions with and without Cre are combined. Schematic of plasmid titration of ZFa.

(B) Output reporter mGL geometric MFI versus input ZFa geometric MFI (proxied by co-expressed TagBFP) for ZFa titration. Values are normalized to the condition without Cre with 0.125X (14 ng of ZFa). Overlaid lines represent model fits with 95% confidence interval. The output shows increasing at low ZFa levels where the distribution is biomodal, followed by a dosage invariant regime.

(C) Single-cell distributions of output mGL expressed from the 203 bp spacer DIAL promoter titrated with ZFa (VP16-ZF43) on plasmids transfected into HEK293T cells. Gate drawn (mGL-A>200) to isolate populations with high output levels of the DIAL promoter. In the presence of Cre, the output mGL increases. At low levels of ZFa, output is bimodal.

(D) Output reporter mGL geometric MFI (gated+ by the line in C) versus input ZFa geometric MFI (proxied by co-expressed TagBFP, as shown in A not gated+ for mGL) in transfected HEK293T cells with the ZFa VP16-ZF43 plasmid titration. Values are normalized to the condition without Cre with 0.125X (14 ng of ZFa). Once gated, the reporter output is dosage invariant throughout the ZFa plasmid titration.

(E) Output reporter mGL geometric MFI of reporter expressing cells from the ZFa (VP16-ZF43) titration (gated by the line in C) versus fraction of cells expressing mGL (above gate in C) transfected in HEK293T cells as described in C. The output geometric MFI is dosage invariant to ZFa plasmid titration. The fraction above the mGL-gate correlates with ZFa level. The fraction has no effect on output level of the cells expressing mGL from the DIAL promoter.

(F) Input TagBFP (proxy for ZFa VP16-ZF43) single-cell distributions as encoded with promoters of varying strengths on plasmids in transfected HEK293T cells. Conditions with and without Cre are combined.

(G) Output reporter mGL geometric MFI versus input ZFa geometric MFI (proxied by co-expressed TagBFP) from different strength promoters with ZFa VP16-ZF43 in transfected HEK293T cells. Values are normalized to the condition without Cre with EF1a promoter. Colored according to legend in F.

(H) Single-cell distributions of output mGL from ZFa with different promoters in transfection of HEK293T cells, as described in G, and colored according to legend in F.

(I) Schematic of two-state transcriptional model of promoter activation. Binding of the ZFa and transcriptional activation is is modeled as a single step. The binding affinity, $K_M$, of ZFa does not change upon editing whereas the putative rate of transcriptional activation, $k_{cat}$, increases upon editing. Full reactions for fitting ZFa plasmid titration data described in

(J) Model parameters after fitting ZFa plasmid titration data for ZFas VP16-ZF37 and VP16-ZF43, as shown in Supplementary Table 3 and 4. Model is described in the Method.

All units for output MFI are arbitrary units (a.u.), and fold change is unitless. Large markers represent the mean of biological replicates with span indicating standard error.

**Figure S8. Single-cell joint distributions for titration of ZFa plasmid.**

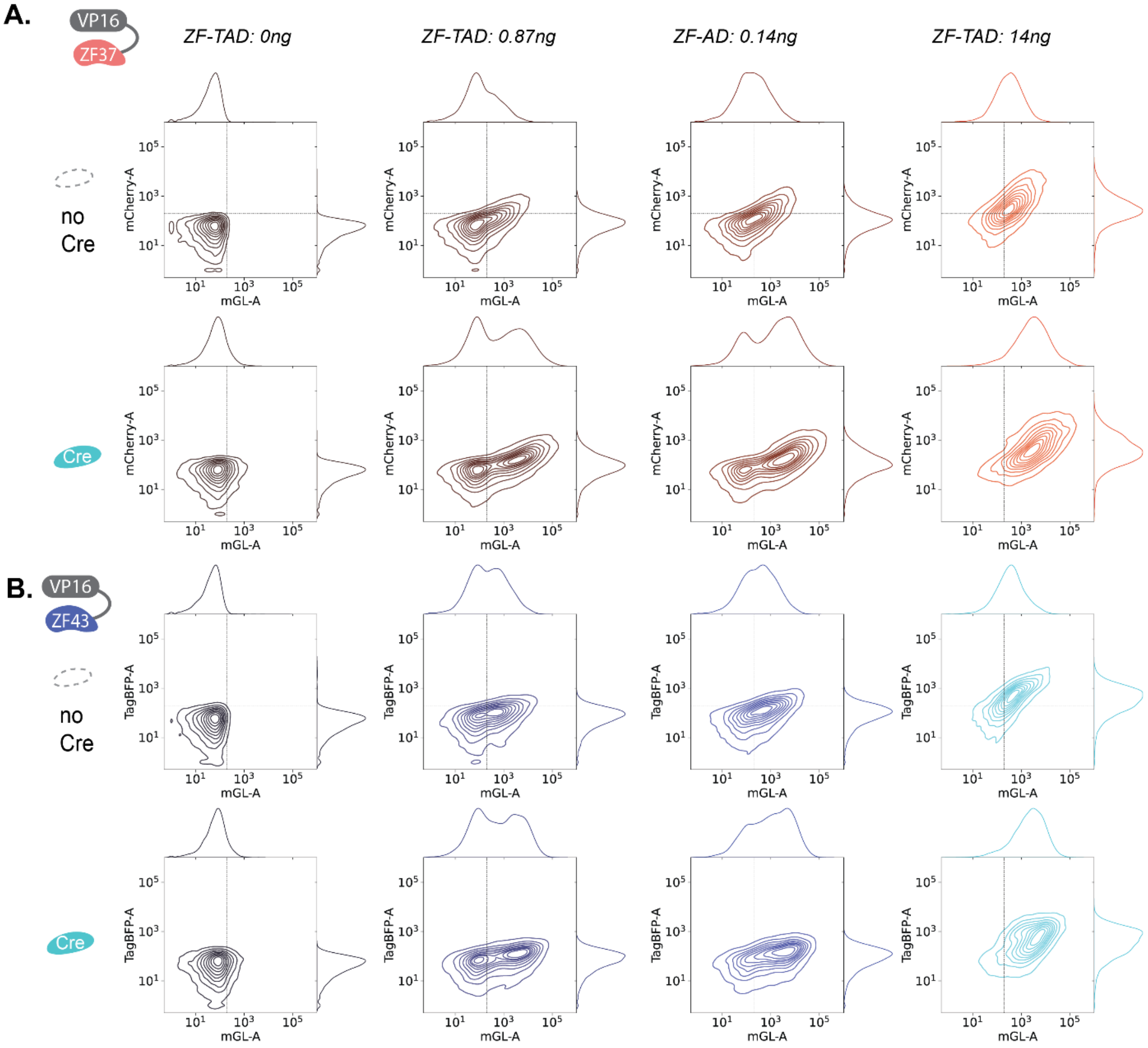


Histograms and joint contour plots of output mGL expressed from the 203 bp DIAL promoter with titrated plasmid amounts of ZF-TAD (A: VP16-ZF37 or B: VP16-ZF43, proxied by co-expressed mCherry or TagBFP respectively) transfected on plasmids into HEK293T cells with or without Cre. Histograms and joint contour plots represent single-cell distributions sample across bioreplicates (n=6 for VP16-ZF37, and n=4 for VP16-ZF43). At low levels of the ZFa (ZF-TAD), output mGL is bimodal. In the bimodal distributions, the cells not expressing the mGL output reporter correspond to the lowest levels of mCherry or TagBFP, suggesting a threshold of ZFa is needed to induce expression from the DIAL promoter.

**Figure S9. DIAL transmits transient inputs into heritable states.**

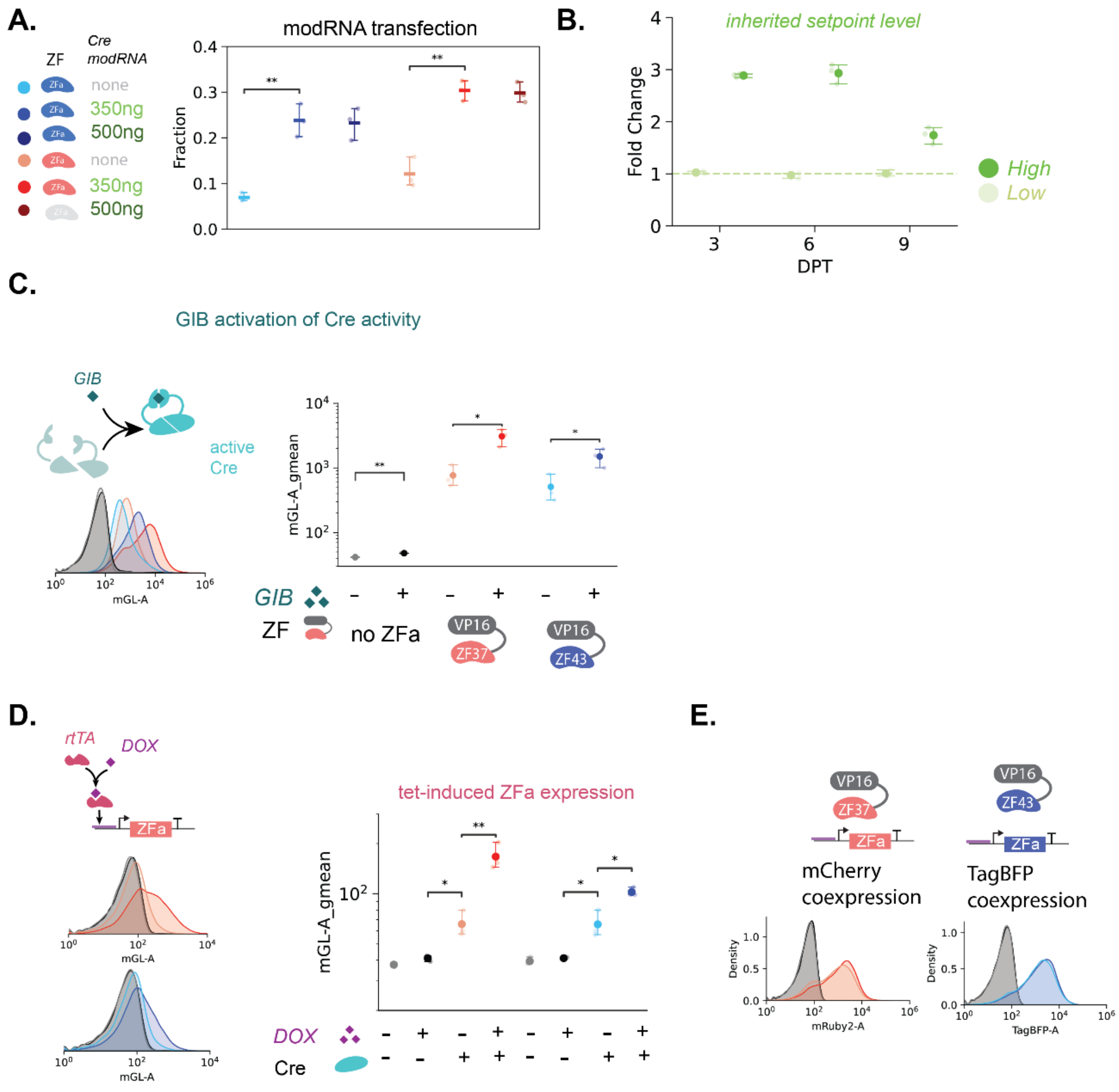


(A) Fraction of output mGL (above a gate based on the condition without Cre) from the 203 bp spacer DIAL promoter with ZFa (VP16-ZF37 or VP16-ZF43) transfected on plasmids into HEK293T cells. Cre ModRNA (0, 350, or 500 ng) was transfected at 1 dpt.

(B) Fold change of the output reporter expressed from the 203 bp spacer DIAL promoter integrated with ZFa into HEK293T cells, co-transfected with Cre modRNA and TagBFP modRNA on day 0, as in Fig 3D. Fold change is the output mGL geometric MFI normalized to condition without Cre at each time point.

(C) Geometric MFI of mGL expressed from the 203 bp DIAL promoter with ZFa and gibberellin-inducible split Cre transfected on plasmids into HEK293T cells. Gibberellin (GIB) induces Cre activity. Histograms depict single-cell distributions of output mGL.

(D) Schematic of doxycycline (DOX) induction of ZFa expression regulated by TRE3G promoter. Geometric MFI of mGL expressed from 203 bp spacer DIAL promoter transfected with TRE3G-ZFa and with or without Cre on plasmids into HEK293T cells. In the plot (right), the first four points are with VP16-ZF37, and the next four points are with VP16-ZF43. Histograms (left) depict single-cell distributions of the output mGL for ZFas, VP16-ZF37 (top) and VP16-ZF43 (bottom), colored according the plot legend.

(E) Single cell distributions of co-expressed mCherry (proxy for VP16-ZF37) or TagBFP (proxy for VP16-ZF43) in D with or without DOX and Cre, colored according to the plot in D.

All units for MFI are arbitrary units (a.u.), and fold change is unitless. Large markers represent the mean of biological replicates with span indicating standard error (n=3). Histograms represent single-cell distributions sampled across bioreplicates (n=3). Statistical significance was calculated with Students t-Test with ns p>0.05; *p<0.05; **p<0.01; *** p<0.001; **** p<0.0001.

**Figure S10. TET-DIAL enables small molecule control of defined setpoints for multiple spacer lengths.**

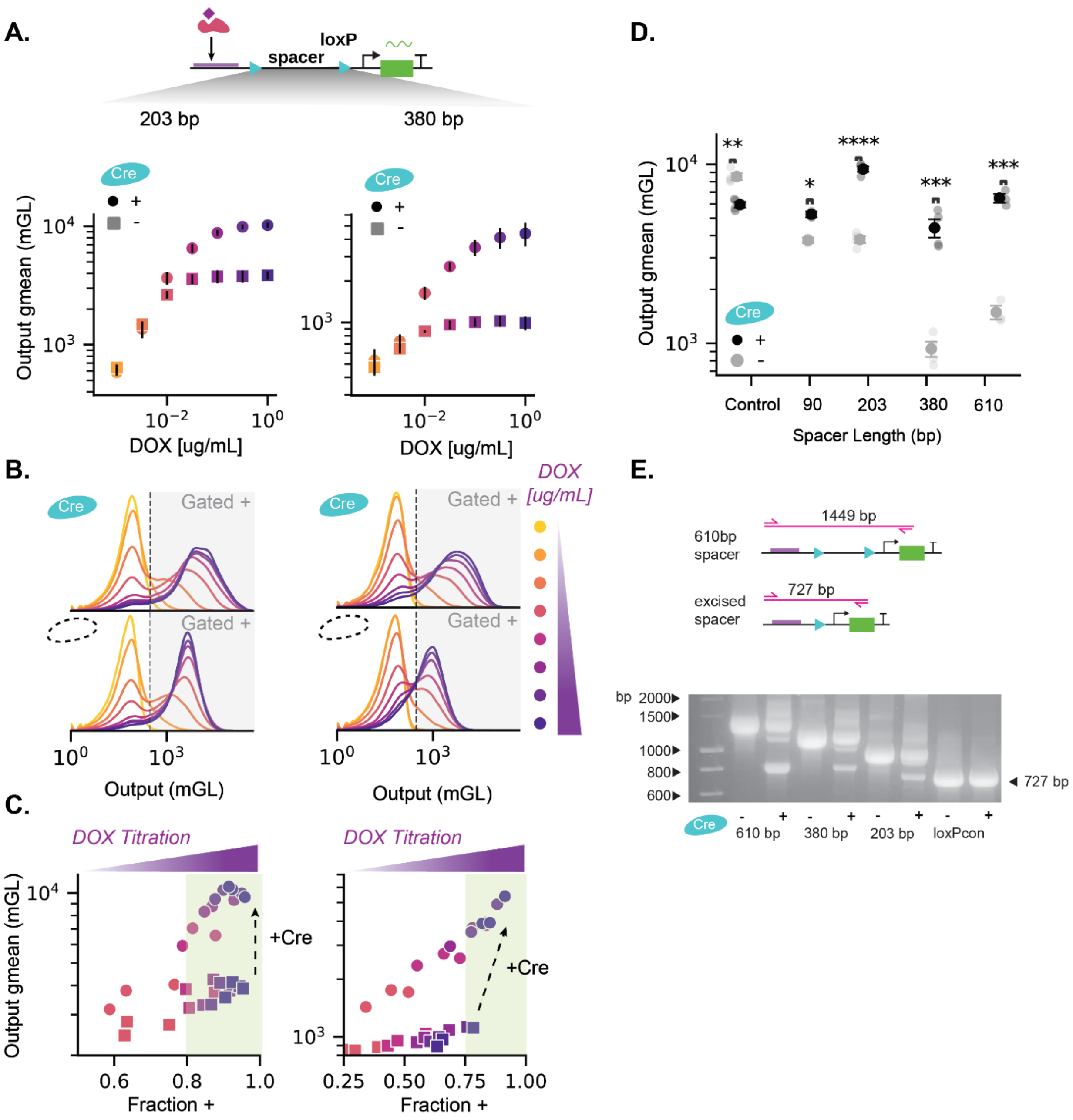


(A) Output reporter mGL geometric mean fluorescence intensity (MFI) from 203 bp or 380 bp spacer TET-DIAL promoter transfected with rtTA on plasmids into HEK293T cells versus input DOX concentration. DOX was induced at 1 dpt.

(B) Single-cell distributions of output mGL expressed from 203 bp or 380 bp spacer TET-DIAL promoter on plasmids transfected with rtTA into HEK293T cells, with varying concentrations of DOX. Colored according to A, with lightest yellow indicating no DOX added. Gates are drawn to isolate cells above the background (no DOX) condition. In the presence of Cre, the output mGL increases. At low concentrations of DOX, output is bimodal.

(C) Output reporter mGL geometric MFI of reporter expressing cells (gated as shown in B) versus fraction of cells expressing mGL (above gate in B) from transfection of HEK293Ts cells as described in B. Colored according to A. Adding Cre induces a new setpoints of reporter expression that can be varied along with fraction of expressing cells by changing DOX concentrations. To set unimodal expression levels (off, low, and high) with high fraction positive in the “ON” state (green region), presence of DOX at high concentration can control whether expression is “OFF” or “ON”, and presence of Cre can control the level of “ON” expression.

(D) Geometric MFI of mGL expressed from TET-DIAL promoters of varying spacer lengths transfected with rtTA on plasmids into HEK293T cells, with or without Cre and with DOX (1 μg/mL).

(E) Schematic of PCR used to visualize changes to the molecular state of the TET-DIAL promoter, labeled with expected PCR product sizes for the 610 bp spacer TET-DIAL promoter. The primers bind to sequences upstream of the TET binding sites or the mGL reporter gene. In the presence of Cre, the spacer excision shortens the distance between binding sites and the minimal promoter, resulting in a shorter PCR product. Image of gel electrophoresis of PCR products from TET-DIAL promoters of various spacer lengths in C. Conditions without Cre are PCR products from the plasmid. Conditions with Cre are PCR products from cell lysis of HEK293T cells transfected with plasmids for Cre, rtTA, and TET-DIAL promoter of corresponding spacer length. The conditions with Cre show a shorter band at 727 bp, the expected length of the PCR product from the post-excision promoter, suggesting the emergence of a post-excision promoter state in the presence of Cre.

All units for MFI are arbitrary units (a.u.). Large markers in (A) and (D) represent the mean of biological replicates (n=3) with span indicating standard error. Markers in (C) represent biological replicates (n=3). Histograms represent single-cell distributions sampled across bioreplicates. Statistical significance was calculated with Students t-Test with ns $p>0.05$; *$p<0.05$; **$p<0.01$; *** $p<0.001$; **** $p<0.001$

**Figure S11. Expression of rtTA influences the bimodality of the TET-DIAL promoter.**

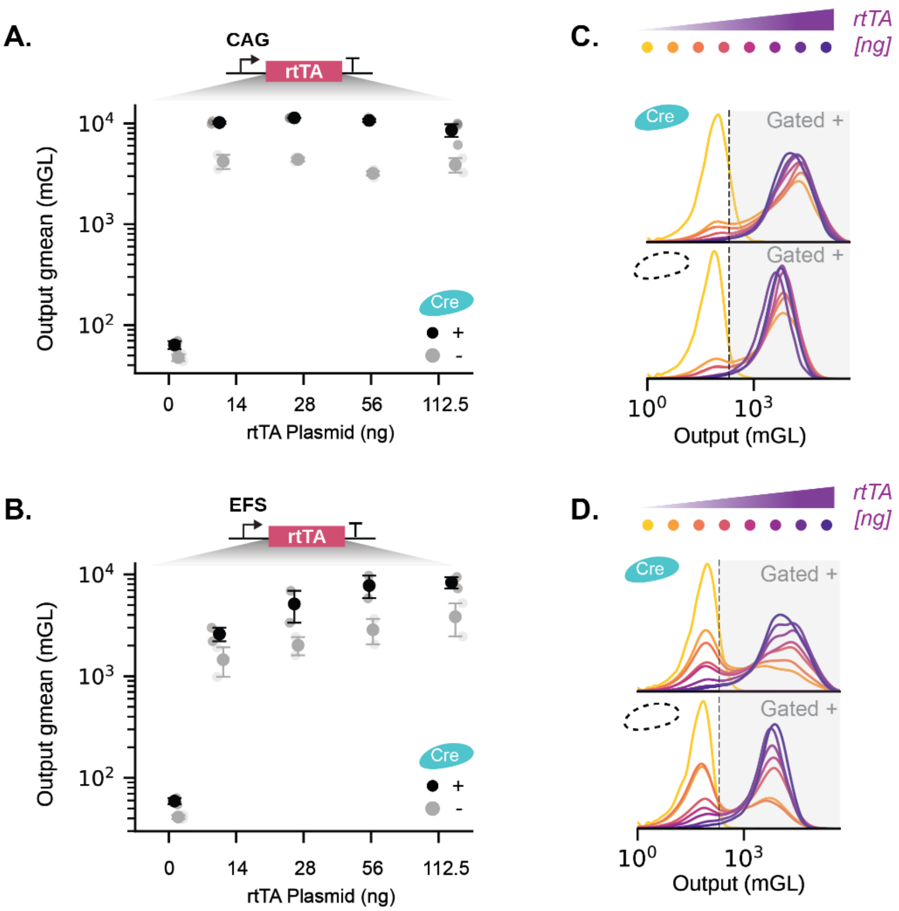


(A-B) Geometric mean fluorescence intensity (MFI) of mGL expressed from 203 bp spacer TET-DIAL promoter with titrated levels of rtTA co-transfected on plasmids into HEK293T cells, with or without Cre. All conditions are at 1x DOX (1 ug/mL). The transactivator rtTA was constitutively expressed from CAG (A) or EFS (B) promoter.

(C-D) Representative histograms for A and B. Single-cell distributions of output mGL expressed from the 203 bp spacer TET-DIAL promoter titrated with CAG-rtTA (C) or EFS-rtTA (D) transfected on plasmids into HEK293T cells, with or without Cre.

All units for MFI are arbitrary units (a.u.). Large markers represent the mean of biological replicates with span indicating standard error. Histograms represent single-cell distributions sampled across bioreplicates (n=3).

**Figure S12. DIAL delivered to MEFs via lentivirus; Cre does not affect the level of ZFa expression.**

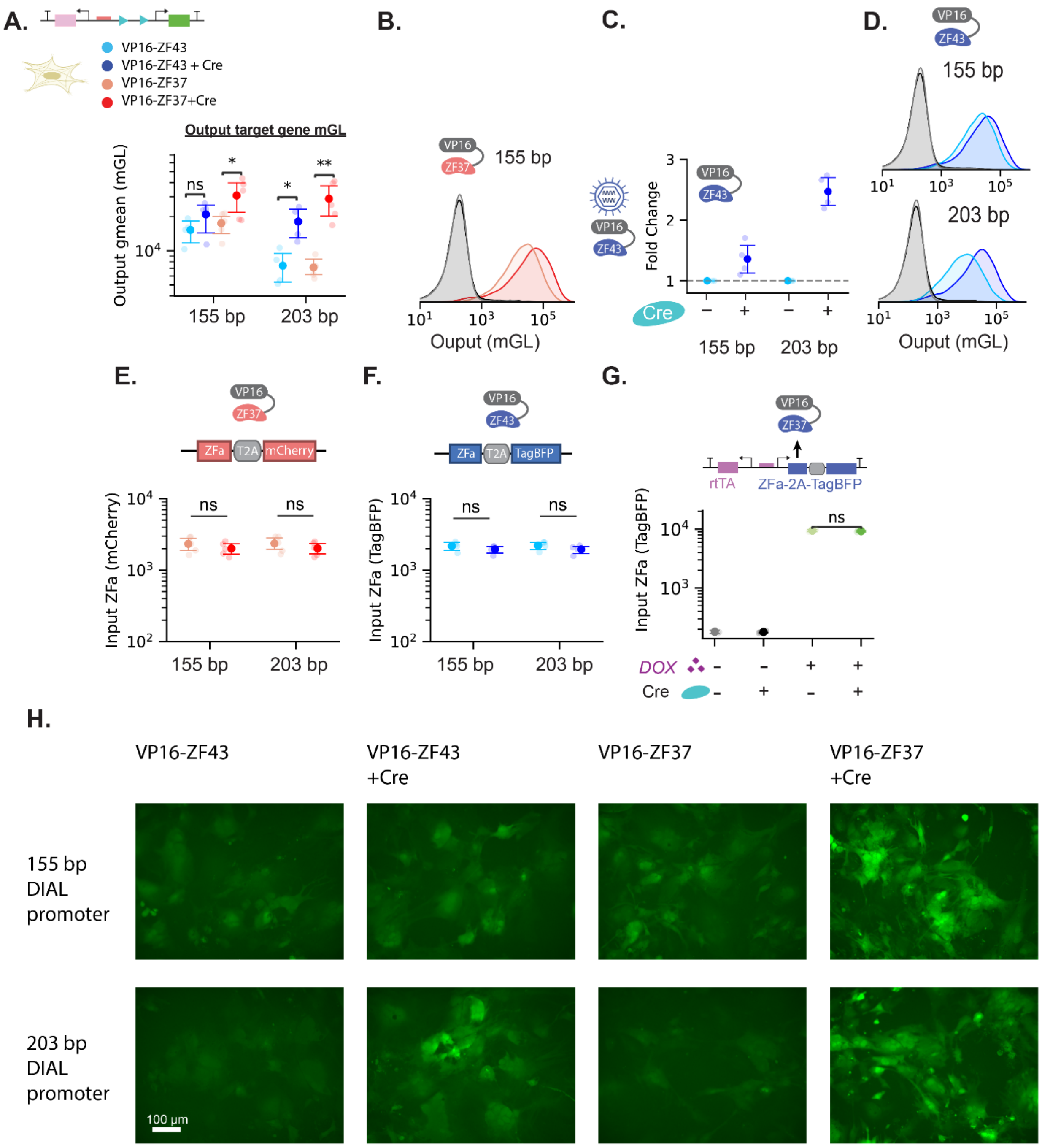


(A) Geometric mean fluorescence intensity (MFI) expressed from 155 bp and 203 bp spacer DIAL promoter delivered via lentivirus with ZFa retrovirus into mouse embryonic fibroblasts (MEFs), with or without Cre retrovirus. Cells were gated for co-delivered iRFP670 expression, co-expressed TagBFP for VP16-ZF43 expression if present, and co-expressed mCherry for VP16-ZF37 expression if present.

(B) Single-cell distributions of output mGL expressed from 155 bp spacer DIAL promoter delivered via lentivirus with or without ZFa retrovirus VP16-ZF37 or Cre. The conditions represent no ZFa and no Cre (gray), no ZFa and Cre (black), VP16-ZF37 and no Cre (light red), and VP16-ZF37 and Cre (dark red). Cells were gated by expression of co-delivered iRFP670. In the presence of ZFa, cells were also gated by expression of mCherry co-expressed with VP16-ZF37.

(C) Fold change of the output mGL reporter expressed from 155 bp and 203 bp spacer DIAL promoters delivered via lentivirus with ZFa retrovirus VP16-ZF43 into MEFs, with or without Cre retrovirus. Fold change is the output geometric MFI normalized to the condition without Cre within each spacer length. Cells were gated by expression of co-delivered iRFP670 and by expression of TagBFP co-expressed with VP16-ZF43.

(D) Single-cell distributions of output mGL expressed from 155 or 203 bp spacer DIAL promoter delivered via lentivirus with or without ZFa retrovirus VP16-ZF43 or Cre. The conditions represent no ZFa and no Cre (gray), no ZFa and Cre (black), VP16-ZF43 and no Cre (light blue), and VP16-ZF43 and Cre (dark blue). Cells were gated by expression of co-delivered iRFP670. In the presence of ZFa, cells were also gated by expression of TagBFP co-expressed with VP16-ZF43.

(E) Geometric MFI of mCherry co-expressed with ZFa VP16-ZF37 delivered via retrovirus with 155 bp or 203 bp spacer DIAL promoter delivered via lentivirus into MEFs, with and without Cre retrovirus. Cells were gated by expression of co-delivered iRFP670 and by expression of mCherry co-expressed with VP16-ZF37.

(F) Geometric MFI of TagBFP co-expressed with ZFa VP16-ZF43 delivered via retrovirus with 155 bp or 203 bp spacer DIAL promoter delivered via lentivirus into MEFs, with and without Cre retrovirus. Cells were gated by expression of co-delivered iRFP670 and by expression of TagBFP co-expressed with VP16-ZF43.

(G) Geometric MFI of TagBFP co-expressed with DOX-inducible ZFa VP16-ZF43 delivered via lentivirus with 203 bp spacer DIAL promoter via lentivirus into MEFs, with or without Cre retrovirus. Adding Cre does not affect DOX-inducible expression of ZFa. Cells were gated by expression of co-delivered iRFP670. In the presence of DOX, cells were also gated by expression of TagBFP co-expressed with VP16-ZF43.

(H) Representative fluorescence microscopy images of mGL expressed from 155 or 203 bp spacer DIAL promoter infected via lentivirus into MEFs with ZFa retrovirus VP16-ZF37 or VP16-ZF43, and with or without Cre retrovirus. Images taken 3 days post-infection (dpi).

All units for MFI are arbitrary units (a.u.), and fold change is unitless. Large markers represent the mean of biological replicates with span indicating standard error. Histograms represent single-cell distributions sampled across bioreplicates (n=3). Statistical significance was calculated with Students t-Test with ns $p>0.05$; *$p<0.05$; **$p<0.01$; *** $p<0.001$; **** $p<0.0001$.

**Figure S13. DIAL is portable to iPSCs; ZFa level is unaffected by recombinase expression.**

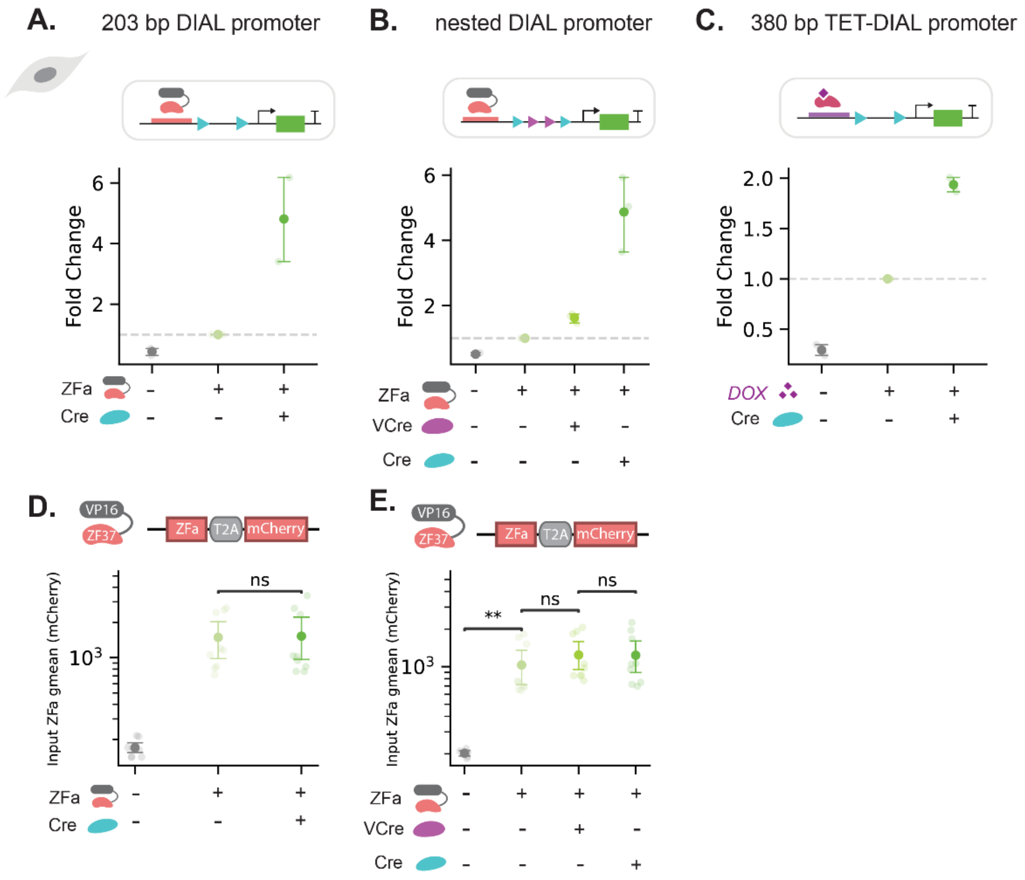


(A, B, C) Fold change of the output reporter expressed from 203 bp spacer DIAL, nested DIAL, and 380 bp spacer TET-DIAL promoters transfected on plasmids into human induced pluripotent stem cells (iPSCs), with and without plasmids for ZFa, VCre, and Cre. Fold change is the output mGL geometric mean fluorescence intensity (MFI) normalized to the condition with promoter and ZFa only.

(D, E) Geometric MFI of mCherry coexpressed with ZFa VP16-ZF37 with 203 bp spacer (shown in D) or nested (shown in E) DIAL promoter transfected on plasmids into iPSCs, with or without plasmids for ZFa, VCre, and Cre. The presence of recombinase does not affect levels of ZFa.

All units for MFI are arbitrary units (a.u.), and fold change is unitless. Large markers represent the mean of biological replicates with span indicating standard error (n=3).Statistical significance was calculated with Students t-Test with ns $p>0.05$; *$p<0.05$; **$p<0.01$; *** $p<0.001$; **** $p<0.0001$.

**Figure S14: Cre does not affect ZFa levels when DIAL promoters regulate different target genes.**

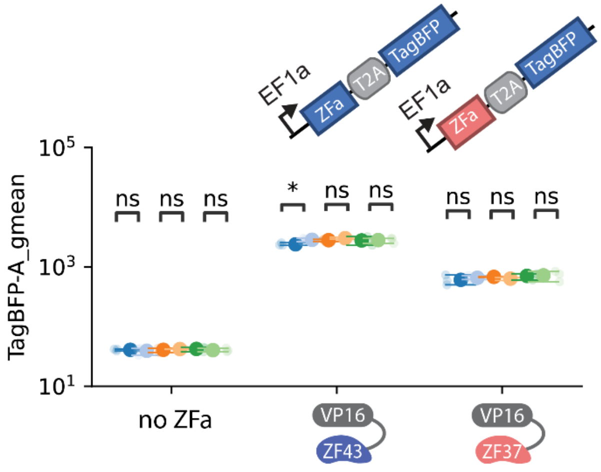


Geometric mean fluorescence intensity (MFI) of input TagBFP (proxy for ZFa VP16-ZF43 or VP16-ZF37) on plasmids transfected into HEK293T cells. Conditions were co-transfected with the 203 bp-spacer DIAL promoter regulating different target genes, and with (light) or without (dark) Cre. Across diverse target genes like mGL-2A-mCherry (blue), Halo-p53 (orange), or mCherry-HRas$^{G12V}$ (green), presence of Cre did not substantially affect ZFa expression levels. All units for MFI are arbitrary units (a.u.). Large markers represent the mean of biological replicates with span indicating standard error (n=3). Statistical significance was calculated with Students t-Test with ns p>0.05; *p<0.05.

**Supplementary Table 1: Plasmid Amounts for Transfection into HEK293T Cells or iPSCs**

| Figure | Description |
|---|---|
| Figure 1<br>Figure S1-S6 | HEK293Ts were transiently transfected with DIAL reporters containing mGL target gene (112.5ng), transfection marker (iRFP670, 112.5ng), with or without ZFa (C,F: 112.5ng; D,E,H,I:14ng), and with or without Cre (11 ng). All conditions were filled with an empty plasmid vector to achieve the same amount of total DNA per condition (349 ng). |
| Figure 2,<br>Figure S7-S8 | HEK293Ts were transiently transfected with the 203bp-spacer promoter with mGL target gene (112.5ng), transfection marker (iRFP670, 112.5ng), with or without Cre (11 ng), and ZF-TAD (2:1 plasmid titration: 0 to 112 ng, different promoters: 112 ng). All conditions were filled with an empty plasmid vector to achieve the same amount of total DNA per condition (349 ng). |
| Figure 3F<br>Figure S9C | HEK293Ts were transiently transfected with the 203bp-spacer promoter with mGL target gene (112.5ng), transfection control (iRFP670, 112.5ng), each half of split GIB-inducible Cre (56ng each), and with or without ZFa or empty plasmid vector (112.5ng), for a total of 450ng DNA per condition. At 1 dpt GIB (1µM) was added. |
| Figure 3G<br>Figure S9D-E | HEK293Ts were transiently transfected with the 203bp-spacer DIAL mGL reporter (112.5ng), transfection marker (iRFP670, 112.5ng), TRE3G-VP16-ZF37-2A-mCherry-BGH or TRE3G-VP16-ZF43-2A-TagBFP-BGH (14ng), rtTA (28ng), and with or without Cre (11ng). All conditions were filled with an empty plasmid vector to achieve same total DNA per condition (349 ng). At 1 dpt, DOX (1 ug/mL) was added. |
| Figure S9A | HEK293Ts were transiently transfected with DIAL reporters containing mGL target gene (112.5ng), transfection marker (iRFP670, 112.5ng), and with or without ZFa (C, F: 112.5ng; D,E,H,I:14ng. All conditions were filled with an empty plasmid vector to achieve the same amount of total DNA per condition (349 ng). ModRNA Cre was delivered at 1 dpt. |
| Figure 4<br>Figure S10 | HEK293Ts were transiently transfected with the TET-DIAL promoter regulating the mGL target gene at varying spacer lengths (112.5 ng), transfection marker (iRFP670, 112.5ng), with or without Cre (11 ng), and EFS-rtTA-2A-mRuby2 (112.5 ng). All conditions were filled with an empty plasmid vector to achieve the same amount of total DNA per condition (349 ng). |
| Figure S11 | HEK293Ts were transiently transfected with the 203 bp-spacer TET-DIAL promoter regulating the mGL target gene (112.5 ng), transfection marker (iRFP670, 112.5ng), with or without Cre (11 ng), and either EFS-rtTA-2A-mRuby2 or CAG-rtTA-2A-TagBFP (2:1 plasmid titration: 0 to 112.5 ng). All conditions were filled with an empty plasmid vector to achieve the same amount of total DNA per condition (349 ng). |
| Figure 5E-H<br>Figure S13 | iPSCs were transiently transfected with DIAL promoters regulating the mGL target gene (203 bp, nested, or TET-DIAL; 100 ng), transfected marker (TagBFP, 25 ng), with or without rtTA or ZFa (12.5 ng), and with or without Cre or VCre (25 ng). All conditions were filled with an empty plasmid vector to achieve the same amount of total DNA per condition (162.5 ng). |
| Figure 5I-K<br>Figure S14 | HEK293Ts were transiently transfected with 203bp DIAL reporter with various target genes (112 ng), ZFa (VP16-ZF37-2A-TagBFP, or VP16-ZF43-2A-TagBFP) (112.5 ng), transfection marker (iRFP670, 112.5ng), with or without Cre (11 ng), and ZF-TAD (112 ng). |

**Supplementary Table 2: Instrument specifications of analytical flow cytometry**

| Experiments | Fluorescent Protein | Excitation Laser (nm) | Laser Setting (V) | Emission Filter (nm) |
|---|---|---|---|---|
| **Figure 1-3** | | FSC | 60 | |
| **Figure 5I-K** | | SSC | 360 | |
| **Figure S1-S9** | mGreenLantern | Blue – 488 nm | 220 | BL1 (510/10) |
| **Figure S14** | Halo (Janelia Fluor 549) | Yellow – 561 nm | 390 | YL1 (585/16) |
| | mCherry | Yellow – 561 nm | 296 | YL2 (615/25) |
| | TagBFP | Violet – 405 nm | 190 | VL1 (440/50) |
| | iRFP670 | Red – 637 nm | 280 | RL1 (670/14) |
| **Figure 4** | | FSC | 60 | |
| **Figure S10-S11** | | SSC | 360 | |
| | mGreenLantern | Blue – 488 nm | 220 | BL1 (510/10) |
| | mCherry | Yellow – 561 nm | 295 | YL2 (615/25) |
| | TagBFP | Violet – 205 nm | 190 | VL1 (440/50) |
| | iRFP670 | Red – 637 nm | 280 | RL1 (670/14) |
| **Figure 5E-H** | | FSC | 60 | |
| | | SSC | 340 | |
| | mGreenLantern | Blue – 488 nm | 380 | BL1 (510/10) |
| | mCherry | Yellow – 561 nm | 440 | YL2 (615/25) |
| | TagBFP | Violet – 405 nm | 300 | VL1 (440/50) |
| | iRFP670 | Red – 637 nm | 460 | RL1 (670/14) |
| **Figure 5B-D** | | FSC | 60 | |
| **Figure S12A-F, H** | | SSC | 345 | |
| | mGreenLantern | Blue – 488 nm | 260 | BL1 (510/10) |
| | mCherry | Yellow – 561 nm | 300 | YL2 (615/25) |
| | TagBFP | Violet – 405 nm | 210 | VL1 (440/50) |
| | iRFP670 | Red – 637 nm | 270 | RL1 (670/14) |
| **Figure 5E** | | FSC | 60 | |
| **Figure S12G** | | SSC | 325 | |
| | mGreenLantern | Blue – 488 nm | 250 | BL1 (510/10) |
| | mCherry | Yellow – 561 nm | 320 | YL2 (615/25) |
| | TagBFP | Violet – 405 nm | 220 | VL1 (440/50) |
| | iRFP670 | Red – 637 nm | 250 | RL1 (670/14) |

**Supplementary Table 3: VP16-ZF37 Plasmid Titration Fitted Model Parameters**

| Parameters | No Cre | Cre |
|---|---|---|
| $k'_{cat}$ | 0.99 | 15.63 |
| $K_M$ | 0.21 | 0.52 |
| $\alpha'$ | 0.14 | 0.22 |

**Supplementary Table 4: VP16-ZF43 Plasmid Titration Fitted Model Parameters**

| Parameters | No Cre | Cre |
|---|---|---|
| $k'_{cat}$ | 1.13 | 5.21 |
| $K_M$ | 0.19 | 0.20 |
| $\alpha'$ | 0.08 | 0.08 |
